# PhenoBIC: operator-free single-cell spatial phenotyping in multiplex imaging data using deep learning of cell staining patterns

**DOI:** 10.64898/2026.06.11.731702

**Authors:** Abishek Sankaranarayanan, Chenkai Zhao, Madeline Gabriela Hernandez, Elizabeth A. Clemens, Kimberly S. Smythe, Anum S. Kazerouni, Lisa L. Carr, Christopher I. Li, Savannah C. Partridge, Shaveta Vinayak, Shachi Mittal

## Abstract

Multiplex imaging is a valuable tool for spatially examining tissue microenvironments at the single-cell level to uncover biological and clinical insights. However, most multiplex image analysis workflows currently require manual intervention for cell phenotyping, which slows progress, demands human effort, and yields operator-dependent outputs. Here, we developed PhenoBIC, a pre-trained deep learning model for image classification of the multiplexed biomarker signals in a cell (**<u>B</u>**iomarker **I**mprint of a **<u>C</u>**ell) to classify cell phenotypes. We show that PhenoBIC (F1-score ∼0.88) outperforms manual gating (widely used) and other machine learning-based computational approaches for cell marker expression classification. We validated this across multiple biomarkers, tissue sampling strategies (whole biopsies and tissue microarrays), multiplex panels, imaging platforms, and tissue types. We have released our in-house training and validation datasets of ∼1.4 million manually curated cell expression ground truth labels. We have also open-sourced PhenoBIC and enabled its community-wide deployment via the QuPath interface.

## Introduction

Multiplex imaging is increasingly being used for single-cell spatial profiling of tissue microenvironments to generate fundamental biological and clinically relevant insights^1–6^. A single multiplex image can profile dozens of proteins across millions of cells and cannot be fully utilized with manual analysis. Image analysis workflows are needed to parse the extensive data in digital multiplex images and generate human-interpretable insights^7^.

In a typical single-cell multiplex image analysis workflow, cell segmentation is used to delineate individual cells in a tissue sample. Although this was a manually laborious process requiring human operators to optimize sample-specific parameters in classical algorithms^8^, such as Otsu thresholding^9^ and watershed^10^, the emergence of pre-trained deep learning models has made segmentation automated, less subjective, and more accurate than operator-driven workflows^11–19^.

The next step is cell phenotyping, where each cell is assigned a phenotype based on its co-expression of biomarkers in the multiplex panel. However, in contrast to segmentation, cell phenotyping still predominantly remains manual and operator-dependent. The commonly used method for estimating cell biomarker expression is manual marker intensity thresholding (“gating”)^20–22^. If the mean intensity signal for a biomarker within the cell area is above this threshold, the cell is “positive” for that marker; otherwise, it is “negative”. Manual gating presents some disadvantages: it is exceedingly labor-intensive to perform gating on each sample and each channel individually, and it is inherently subjective to the operator. Also, there exists an inherent limitation in reducing marker expression to cell averages (integrated expression), as this method does not account for subtle variations in staining patterns within and across cells (e.g., localization to the nucleus/membrane, staining completeness).

Over the past decade, machine learning-based tools have been developed^23–32^ to address these limitations, especially for higher-plex panels. Tools such as FlowSOM^25^ and PhenoGraph^23^ improved throughput but retained dependence on integrated expressions, which do not account for subcellular staining patterns. More recent supervised approaches such as STELLAR^26^ and MAPS^27^ reduced the need for manual annotations but still required dataset-specific retraining with limited generalizability across diverse marker panels and imaging platforms. These tools still either rely on cell marker intensity averages^23–30^ and/or require user intervention^23–27,31,32^ in the form of cluster manipulation/labelling or dataset-specific training annotations.

Similar to the trajectory of cell segmentation, pre-trained deep learning models that directly read multiplex image data have the potential to address these limitations and fully automate robust cell phenotyping. Other groups have recently trained deep learning models to push the field in this direction^33–36^, highlighting both the promise of and the need for methods that reliably scale across heterogeneous imaging contexts. Further advances are needed to overcome key challenges in achieving fully automated cell phenotyping both with robust performance and computational efficiency, and these tools must be deployed in an accessible, no-code interface for broad community adoption.

Here, we address these gaps by presenting a fully automated deep learning platform, PhenoBIC (https://github.com/Shachi-Mittal-Lab/PhenoBIC). PhenoBIC introduces a cell phenotype classification paradigm centered on the **<u>B</u>**iomarker **I**mprint of a **<u>C</u>**ell. It captures the characteristic spatial staining pattern encoded within a cell region learned through a state-of-the-art lightweight convolutional neural network. Unlike previous approaches that predict cell types directly, PhenoBIC carries out inference independently for each channel to determine marker positivity and assign cell phenotypes based on co-marker positivity combinations. This enables generalizable performance across multiplex panels with any combination of markers without retraining. To support this, we have curated and released (https://huggingface.co/datasets/mittal-research-lab/PhenoBIC-Data) the largest manually annotated, quality-controlled cell expression dataset compared to previous studies, consisting of approximately 1.4 million gold-standard cell marker expression labels. PhenoBIC demonstrates a strong and consistent performance across diverse validation contexts, with a composite F1-score of ∼0.88, illustrating improvement over the current state-of-the art computational phenotyping tool^33^. Furthermore, evaluation of PhenoBIC on external validation datasets comprising independent patient samples, biomarker panels, staining runs, and imaging acquisitions not seen during the training confirms its generalizability and zero-shot learning with strong performance maintained across most markers and patient samples. Beyond just the overall F1-score, PhenoBIC exhibited a substantially more balanced tradeoff between precision and recall. This is particularly crucial for clinical applications as a systematic imbalance between false positives and false negatives can introduce bias in downstream cell frequency estimates, spatial maps, and clinical correlations. We also benchmarked PhenoBIC against manual gating by two independent expert annotators, where it matched or exceeded expert-level classification to deliver objective, operator-free cell phenotyping. Additionally, PhenoBIC significantly reduces computational time compared to similar deep learning-based solutions previously reported, representing a substantial contribution particularly when managing multiplexed samples comprising hundreds of thousands of cells and multiple channels. Additionally, PhenoBIC has been made available as a plug-in (https://github.com/Shachi-Mittal-Lab/PhenoBIC-qupath-extension) deployed within QuPath^37^, an open-source, widely used user interface for visualizing and handling multiplex imaging data, providing the community with a reliable, fully automated phenotyping solution to convert multiplex images to spatial cell maps.

## Results

### Overview of PhenoBIC

PhenoBIC is a pre-trained deep learning tool for cell phenotyping that takes in multiplex imaging data and corresponding cell segmentations as input. For a given multiplex image (Fig. 1A), nuclear segmentation (Fig. 1B) can be conducted utilizing various existing tools, employing a nuclear counterstain channel (Fig. 1C), such as DAPI, Hoechst, etc. Whole cells can be approximated from nuclear segmentations (Fig. 1D), or alternatively, they can be directly segmented using membrane marker channels^11,18^.

**Fig. 1:**
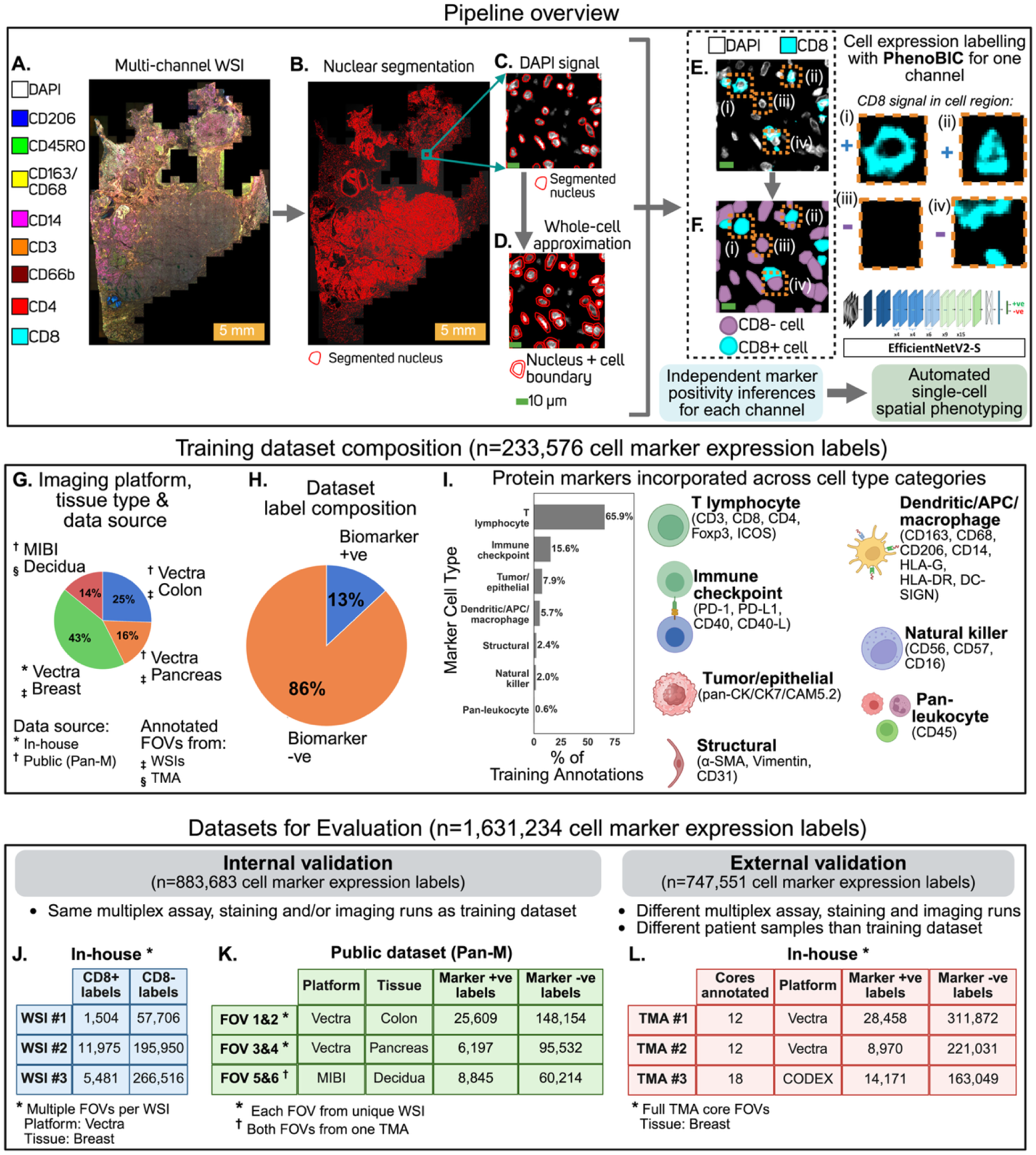
PhenoBIC pipeline and datasets. **A.** Multiplex whole slide image (WSI). **B.** Automated deep learning-based nuclear segmentation using fine-tuned Cellpose model. **C.** Zoomed-in region of interest (ROI) showing DAPI grayscale signal and nuclear segmentations. **D.** Whole-cell segmentation. **E.** DAPI and CD8 channels for zoomed-in ROI. **F.** Classification of CD8 protein expression (i.e., CD8+ vs CD8-) for the cells in the ROI using PhenoBIC- the deep learning model developed in this work with an EfficientNetV2-S architecture. **(i)** and **(ii)** are cropped regions that show the CD8 “imprint” for CD8+ cells, **(iii)** and **(iv)** show the same for CD8- cells. PhenoBIC has learned which imprints correspond to positive vs negative cell expression. **G.** Breakdown of training dataset diversity by imaging platform, tissue type, and data source (in-house vs publicly available). **H.** Breakdown of training labels (positive vs negative for cell marker expression). **I.** Breakdown of biomarker channels from multiplex images. **J.** In-house acquired dataset of multiplex whole slide images (WSIs), with cells in sampled fields of view (FOVs) annotated for biomarker expression. **K.** Annotated FOVs from a public (Pan-M) dataset. J. and K. are considered “internal” validation datasets because they have the same biomarker panel, staining, and/or imaging runs as the training dataset. **L.** “External” validation dataset comprising tissue microarrays (TMAs) of different patient samples, biomarker panels, staining, and imaging runs than the training dataset to evaluate model generalizability.

PhenoBIC is compatible with cell segmentations generated by any upstream tool and is used to classify marker positivity for each cell with respect to each channel. Each cell is cropped to create its own patch for a specific marker channel, acting as an “imprint” of that cell. Examples of cell crops with CD8 cell imprints for CD8+ cells are displayed in Fig. 1E(i) and (ii), while those for CD8-cells are shown in Fig. 1E(iii) and (iv). PhenoBIC has been trained to accurately identify the imprints corresponding to marker-positive and marker-negative cases (Fig. 1F), generating independent per-channel inferences.

### Curating the PhenoBIC datasets

Training and rigorously evaluating a supervised deep learning model for cell expression classification requires large quantities of multiplex images with high confidence and fidelity. To do this, we acquired multiplex images and manually annotated one of the largest cell expression datasets with ∼1.4 million cell marker expression labels. We conducted multiple rounds of quality-controlled reviews (QC) with different expert human annotators to ensure a high standard of training labels (Fig. 2, see Methods).

**Fig. 2:**
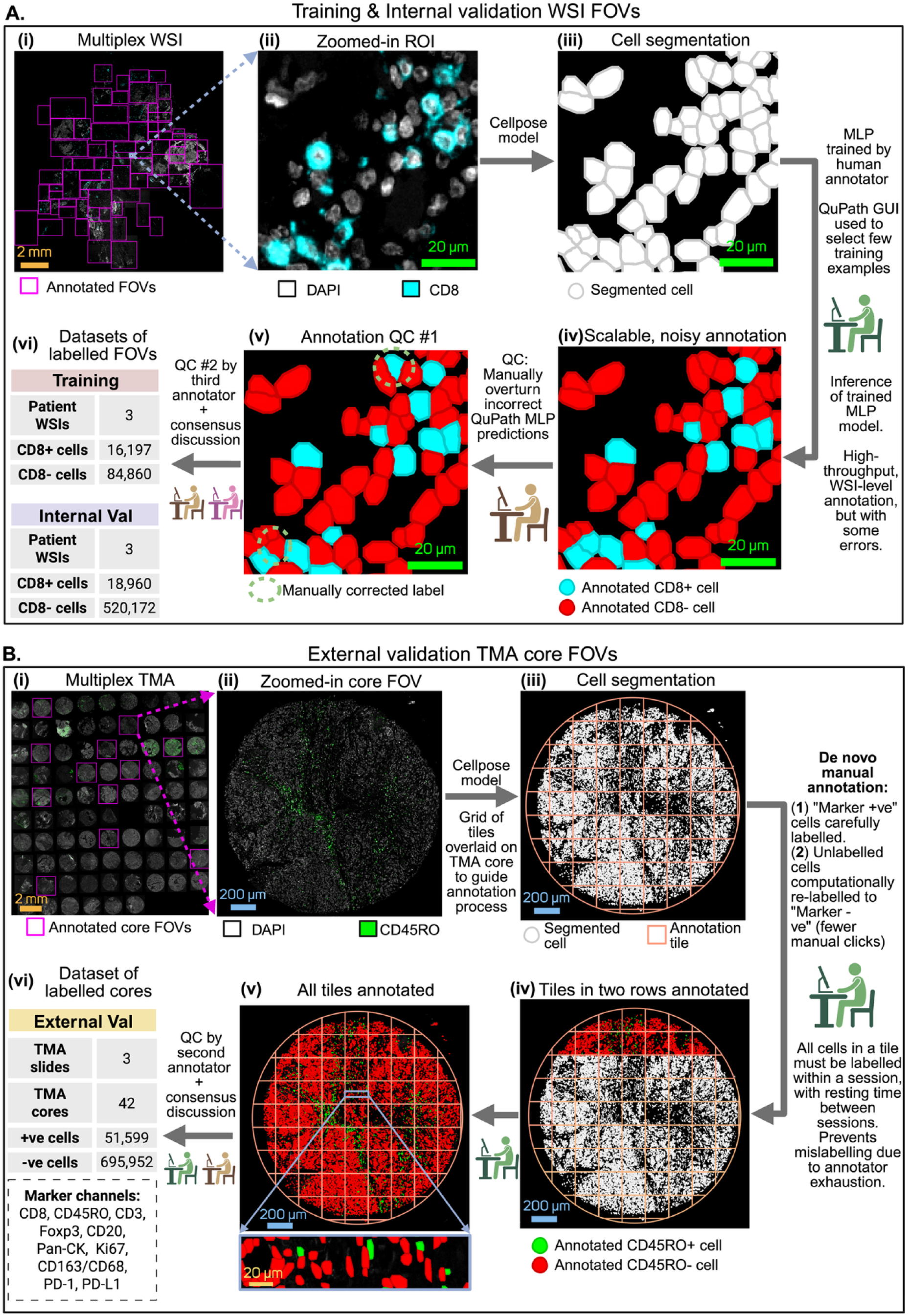
Annotation of in-house datasets. **A.** Annotation of internal validation WSI FOVs. **(i)** Multiplex whole slide image (WSI) with fields of view (FOVs) sampled for annotation. **(ii)** Zoomed-in region of interest (ROI) showing the DAPI and CD8 channels. **(iii)** Nuclear segmentation with a fine-tuned Cellpose model followed by buffering to approximate whole cell segmentations. **(iv)** Multilayer perceptron (MLP-simple machine learning model) trained on CD8 signal statistics (mean, median, maximum, minimum, standard deviation) of each cell to classify cells as CD8+ or CD8-. **(v)** A second human annotator performed quality control (QC) checks and manually overturned incorrect labels by MLP. **(vi)** A third human annotator performed a final QC and discussed potential corrections with the second human annotator to generate consensus labels for ambiguous cases. The in-house training and internal validation datasets were curated in this manner. The number of WSIs and annotated cell marker expression labels are shown for each. **B.** Annotation of external validation TMA core FOVs. **(i)** Multiplex tissue microarray (TMA) with TMA core FOVs sampled for annotation. **(ii)** Zoomed-in FOV of a sampled core showing the DAPI and CD45RO channels. **(iii)** Nuclear segmentation with a fine-tuned Cellpose model followed by buffering to approximate whole cell segmentations. A grid of tiles was overlaid on the TMA core to guide the annotation process. **(iv)** Manual cell labelling (e.g., CD45RO+ vs CD45RO-) by a human annotator. Tile-by-tile progress with scheduled breaks between sessions to prevent mislabelling caused by annotator exhaustion. **(v)** Full TMA core annotated with zoom-in included for clarity. **(vi)** QC check by a second human annotator, who discussed potential label corrections required with the first human annotator to reach a consensus. The external validation dataset was curated in this manner to evaluate model generalizability to biomarker panels, and staining and imaging runs not seen during the training. The number of TMA slides, TMA cores, and annotated cell marker expression labels are shown.

The in-house training and internal validation datasets shared identical biomarker panels, staining procedures, and imaging acquisitions, and may exhibit shared batch effects. Performance evaluations on this internal validation dataset reflect the model’s ability to accurately classify cell marker expression within a controlled and consistent imaging context, providing a reliable upper-bound estimate of the classification accuracy before evaluating generalization to the independent external dataset.

We generated ∼640k cell examples in the in-house training and internal validation datasets (Fig. 2A(vi)), which were labeled for the expression status of a relatively “well-behaved” protein marker, CD8, motivated by the following three properties to enable the learning of robust biomarker imprints. First, there is a significant difference in the quantitative expression of the CD8 protein between the cells designated as CD8 “positive” and “negative”^38^. Second, digital automated counting of CD8+ cells has demonstrated a strong correlation with manual counting of CD8+ cells, validating its reliability in multiplex workflows^39^. Third, the distribution of CD8+ cells is notably homogeneous within the breast cancer tissue sections^40^. This means that we can sample CD8+ cells for training from various regions of a patient’s tissue section.

To ensure that this model is genuinely generalizable across diverse imaging contexts, it was necessary to incorporate additional biomarkers and tissue types exhibiting different staining patterns. To achieve this, we employed Pan-M^33^, a publicly available dataset comprising manually curated cell positivity labels derived from multiplex imaging data. We utilized three tissue subsets of the Pan-M data (with three labelled FOVs from each across training and internal validation): colon tissue imaged using Vectra (PhenoImager™ HT imaging platform), pancreas tissue imaged using Vectra, and decidua tissue imaged using multiplexed ion beam imaging (MIBI) (Fig. 1G).

13% of the composite training dataset was marker-positive labels (Fig. 1H), which is intuitive as typically in tissue, the majority of cell phenotypes exhibit low prevalence. We incorporated a modified loss function during training to prevent the model from defaulting to a low recall result in response to this imbalance. The diversity of the protein markers incorporated in the model training for both marker +ve and –ve expressions and the nature of cell phenotypes they express are depicted in Fig. 1I and Extended Data Fig 1A. Notably, we included heterogeneity from two cell segmentation algorithms, Cellpose and Mesmer, in the training and validation datasets. The quantity of marker expression labels within the internal validation datasets from both the in-house and public data sources is presented in Fig. 1J and 1K, respectively.

To assess the model’s ability to generalize to samples from entirely different patient cohorts, biomarker panels, staining cycles, and imaging runs than those used in training, we acquired and manually annotated (with independent QC) multiple cores across three multiplex breast tissue microarrays (TMAs) (Fig. 2B, see Methods), imaged using Vectra and CODEX (PhenoCycler®) imaging platforms, to serve as the external validation dataset for this study. The number of cell labels annotated across the three TMAs is shown in Fig. 1L, with a detailed breakdown of individual biomarker labels available in Extended Data Fig. 1B.

### PhenoBIC qualitative evaluation

We deployed PhenoBIC on each of the marker channels within our internal and external validation datasets. Fig. 3A(i) displays one of the three multiplex in-house internal validation WSIs, with a representative region of interest (ROI) shown in Fig. 3A(ii) and its associated cell segmentation mask in Fig. 3A(iv). It is evident that the spatial distribution of PhenoBIC’s positive calls (Fig. 3A(v)) closely mirrors the ground truth (Fig. 3A(iii)) annotations, with a spatially resolved color-coded confusion matrix illustrated in Fig. 3A(vi). In this ROI (149 cells), the four false positives resulted from spatial bleed-through of the CD8 signal from adjacent strongly positive cells. Additionally, there is one false negative prediction for a cell that clearly expresses CD8 in a specific and complete manner but appears less bright. Despite these occasional edge cases, PhenoBIC demonstrates robust precision and recall in this ROI.

**Fig. 3:**
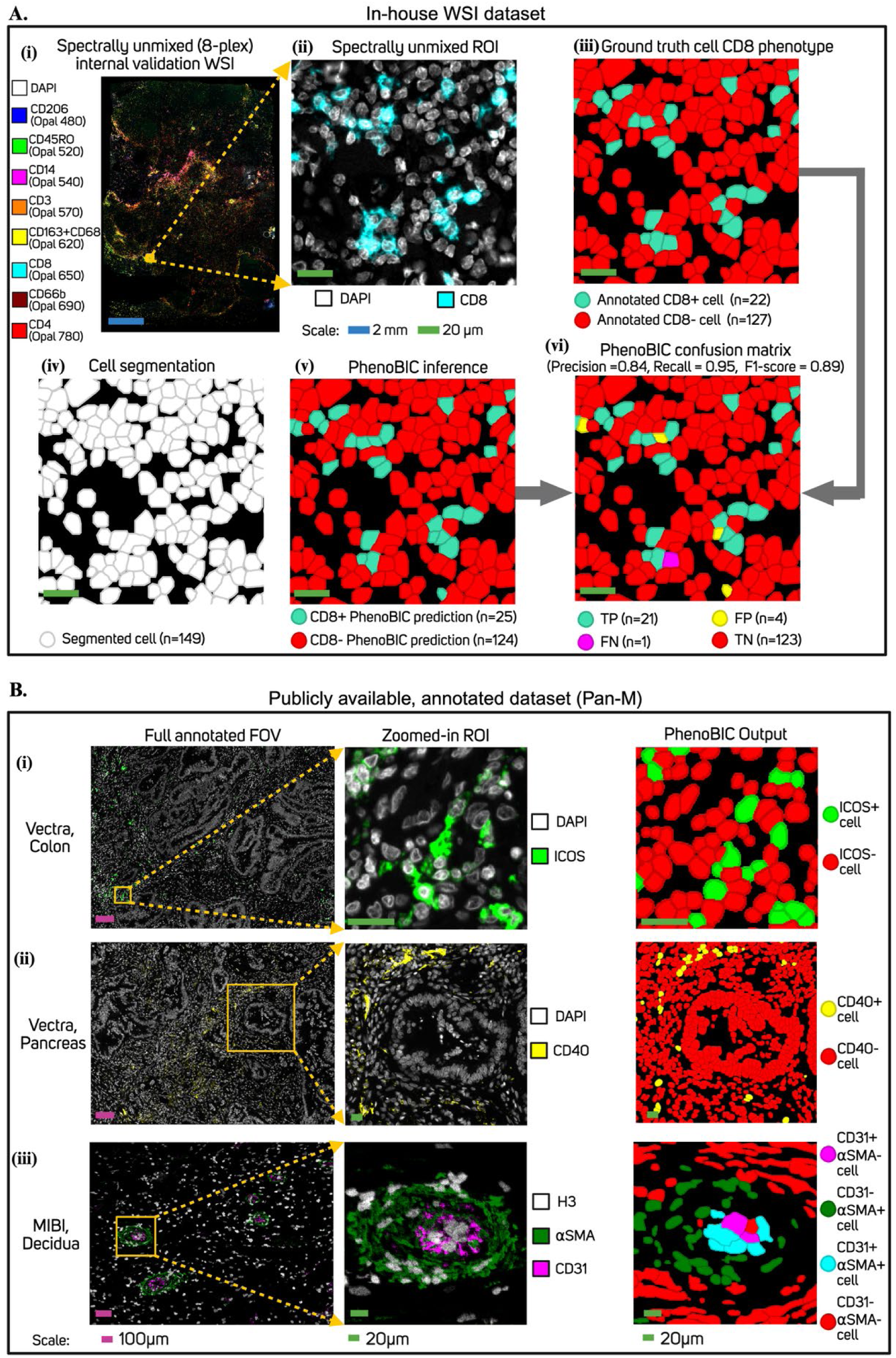
Qualitative evaluation with the internal validation datasets. **A.** In-house whole slide images (WSIs) dataset. **(i)** Representative multiplex WSI. **(ii)** Zoomed-in region of interest (ROI) showing the DAPI and CD8 signal. **(iii)** Ground truth CD8 cell expression labels annotated for the ROI. **(iv)** Cell segmentations for the ROI. **(v)** PhenoBIC prediction of CD8 cell expression. **(vi)** Confusion matrix for the PhenoBIC prediction identifying true positives (TP), false positives (FP), true negatives (TN), and false negatives (FN) for CD8 cell expression. **B.** Publicly available Pan-M dataset with annotated multiplex fields of view (FOVs). The left column shows the multiplex signal of representative annotated FOVs from three sub-datasets: **(i)** colon tissue imaged with Vectra, **(ii)** pancreas tissue imaged with Vectra, and **(iii)** decidua tissue imaged with MIBI. The middle column shows zoomed-in regions of interest (ROIs) with channels of interest. The right column shows the phenotyping output of PhenoBIC.

Fig. 3B demonstrates PhenoBIC’s qualitative performance on the internal validation Pan-M data. PhenoBIC identified ICOS+ activated co-stimulatory T cells within a colon tissue, despite a more diffused staining pattern (Fig. 3B(i)). It correctly outputted CD40+ cells, which are involved in stimulating immune responses, localized to the outer ring of a ductal structure in a pancreas tissue (Fig. 3B(ii)). Fig. 3B(iii) illustrates PhenoBIC’s inference regarding the combinatorial co-expression of CD31 (endothelial marker) and ⍺SMA (vascular smooth muscle) cell positivity in and around a decidual blood vessel.

Fig. 4 A(i), B(i), and C(i) depict the multiplex signals from a core sampled from the external validation TMAs (#1-3, respectively). Fig. 4(ii) shows zoomed-in ROIs from the cores. Fig. 4(iii) illustrates the outputs of PhenoBIC for the entire cores, where each unique cell color corresponds to a co-expression phenotype with biological significance, and Fig. 4(iv) shows the same specifically for the zoomed-in ROIs. Fig 4A(ii) presents a highly immune-infiltrated region with multiple T cell subsets identified by PhenoBIC as shown in Fig. 4A(iv). Fig. 4B(iv) illustrates the profiling of PD-1-expressing cytotoxic T cells, non-expressing counterparts, macrophages, and B cells. PhenoBIC was able to correctly classify four markers with distinct cell type associations and staining patterns in a densely packed multicellular environment. Fig. 4C(iv) depicts the profiling of epithelial/tumor cells and non-epithelial cells, stratified by proliferation status. Ki67 is a marker of cell proliferation with a characteristic nuclear staining pattern, fundamentally different from the membrane-localized markers in the previous two TMAs, indicating PhenoBIC’s ability to effectively handle both localization patterns. Fig. 4(v) indicates the classification confidence enumerated by the number of channels correctly classified by PhenoBIC in the ROIs, determined by comparing the PhenoBIC output to ground truth marker expression labels. PhenoBIC demonstrates robust performance with a low frequency of misclassified channels in the multiplex context, further validated by the quantitative evaluation in the subsequent section.

**Fig. 4:**
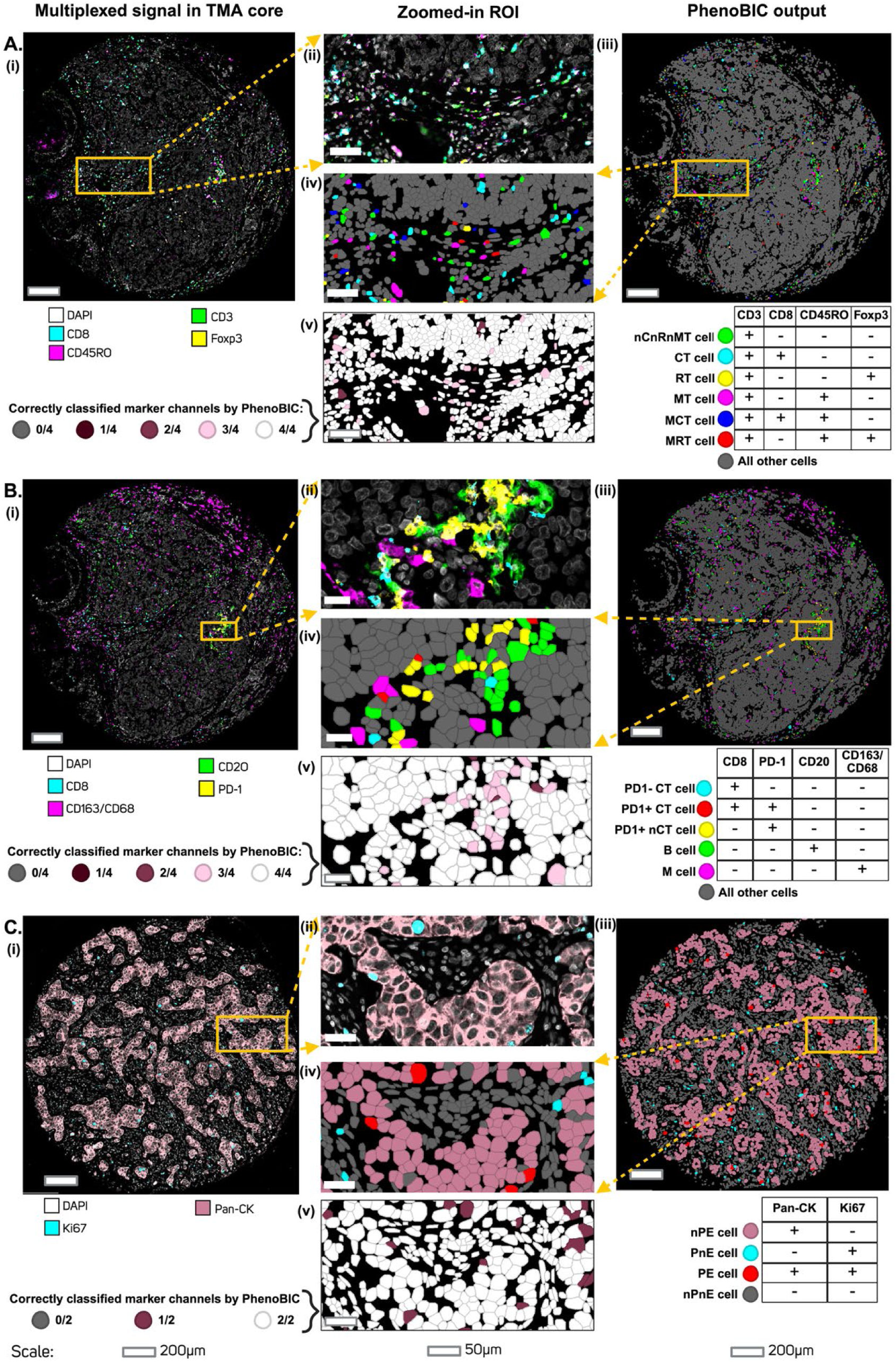
Qualitative evaluation with the external validation dataset. Representative cores are visualized from external validation TMAs **A.** #1, **B.** #2, and **C.** #3. **(i)** Multiplex signal in representative TMA cores with channels of interest shown**. (ii)** Zoomed-in region of interest (ROI) in the multiplex core**. (iii).** Independent PhenoBIC inference for each channel enables cell biomarker co-expression phenotyping. <u>Abbreviations</u> for cell phenotypes are as follows: nCnRnMT cell: non-cytotoxic, non-regulatory, non-memory T cell, CT cell: cytotoxic T cell, RT cell: regulatory T cell, MT cell: memory T cell, MCT cell: memory cytotoxic T cell, MRT cell: memory regulatory T cell, nCT cell: non-cytotoxic T cell, B cell: B cell, M cell: macrophage, nPE: non-proliferating epithelial/tumor cell, PnE: proliferating non-epithelial/tumor cell, PE: proliferating epithelial/tumor cell, nPnE: non-proliferating non-epithelial/tumor cell. **(iv)** Co-expression phenotypes of cells in ROI. **(v)** Accuracy of PhenoBIC in correctly classifying each marker channel expression for the cells in the ROI.

### PhenoBIC quantitative evaluation

Cell marker expression data in multiplex tissue imaging is inherently class-imbalanced, with marker-positive cells typically constituting a small minority of the total cell population. We prioritized the F1-score as our primary evaluation metric as it reflects the model’s ability to correctly identify the biologically meaningful marker-positive population while minimizing both false positives and false negatives, as illustrated by the confusion matrix in Fig. 3A(vi).

For the three in-house WSIs of the internal validation dataset, PhenoBIC achieved a composite F1-score of 0.92 (Extended Data Fig. 2A(i)) and exceeds the performance of manual gating by two independent expert human annotators as well as that of the existing state-of-the-art pre-trained computational approach, Nimbus, in each of the WSIs (Fig. 5C). Across the hold-out validation FOVs in the Pan-M data, PhenoBIC achieved a composite F1-score of 0.89 (Extended Data Fig. 2A(ii)), matching that of Nimbus as reported in their publication^33^. Fig. 5D lists the F1-score of PhenoBIC for individual annotated marker channels across the three Pan-M data subsets, showing strong performance across marker and tissue types. There were some cases where PhenoBIC underperformed, such as with ICOS and PD-L1 in the Vectra Colon dataset, PD-1 in the Vectra Pancreas dataset, and CD57 in the MIBI Decidua dataset, predominantly due to unusually high levels of diffuse staining, further elaborated in the Discussion section.

**Fig. 5:**
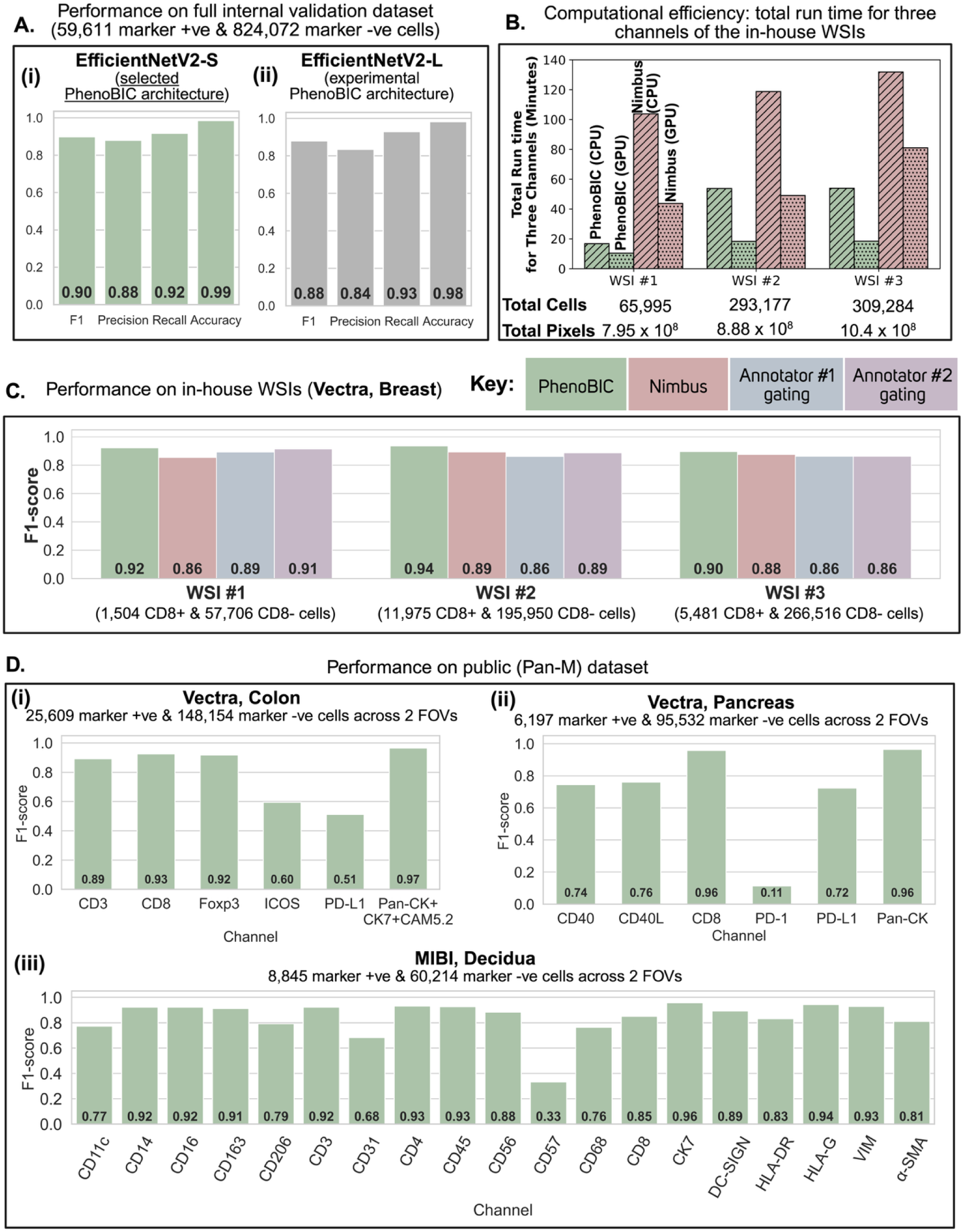
Quantitative evaluation of classifier performance and efficiency using the internal validation dataset. **A.** Evaluation of **(i)** PhenoBIC model (EfficientNetV2-S architecture) and **(ii)** a model trained using another experimental architecture (EfficientNetV2-L), using F1-score, precision, recall, and accuracy metrics. **B.** Evaluation of computational efficiency using total time taken to generate marker positivity calls for three channels of the in-house internal validation WSIs for PhenoBIC and Nimbus using CPU-only and GPU-enabled systems. **C.** CD8 cell expression classification F1-scores for the in-house dataset of PhenoBIC, Nimbus, and manual gating by two expert human annotators. **D.** PhenoBIC F1-scores for various biomarkers in the public (Pan-M) dataset. There were three subsets of the data with two annotated FOVs in each: **(i)** colon tissue imaged with Vectra, **(ii)** pancreas tissue imaged with Vectra, and **(iii)** decidua tissue imaged with MIBI.

PhenoBIC demonstrated strong zero-shot performance on the external validation dataset, which comprised independent patient samples, biomarkers, staining runs, and imaging acquisitions entirely absent from the training data, with a composite F1-score of 0.88 (Fig. 6A(i)). The minimal drop of 0.02 in F1-score relative to the internal validation performance (Fig. 5A(i)) indicates robust generalizability.

**Fig. 6:**
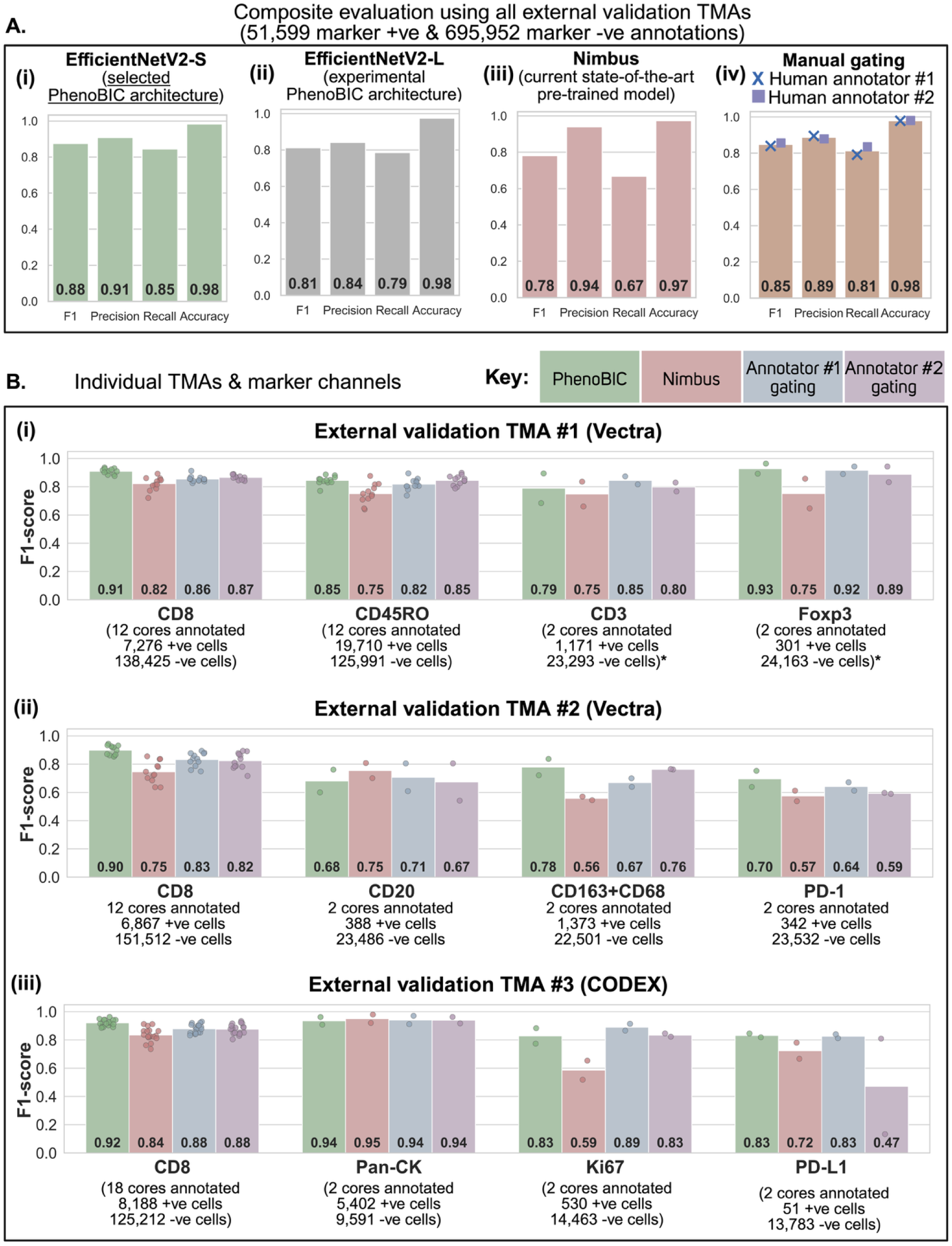
Quantitative evaluation with the external validation dataset. A. Evaluation of **(i)** PhenoBIC model (EfficientNetV2-S architecture), **(ii)** a model trained using another experimental architecture (EfficientNetV2-L), **(iii)** Nimbus- a pre-trained state-of-the-art deep learning model for cell expression classification, and **(iv)** manual gating by two expert human annotators, using F1-score, precision, recall, and accuracy metrics. For manual gating, average cell signal intensity thresholds for marker positivity were visually determined for each channel and each core. **B.** Marker cell expression classification F1-scores of PhenoBIC, Nimbus, and manual gating for the individual external validation TMAs and marker channels annotated. Data points show the F1-scores for individual annotated TMA cores, with the average shown by the bars.

Consistent with the internal validation WSIs, PhenoBIC (Fig. 6A(i)) outperformed both Nimbus (Fig. 6A(iii)) and manual gating by two independent expert human annotators (Fig. 6A(iv)). Again, PhenoBIC demonstrates stable and dependable performance across multiple TMA cores within a slide, different TMA slides, and marker channels, as shown in Fig. 6B. It should be noted that CODEX imaging data (Fig. 6B(iii)) was not seen in the training process, nor were the markers CD45RO, CD20, and Ki67 (Fig. 6B(i)-(iii)). PhenoBIC’s strong performance in these scenarios provides confidence that it will effectively generalize to other imaging platforms and biomarkers not included within this study.

It is also important to recognize that the manual gating here represents the best-case scenario. Both annotators visually optimized the best threshold for marker positivity gating of each WSI/TMA core separately, ensuring robust performance across patient samples. However, in practice it is too time-consuming to optimize the thresholds for each WSI/TMA core. Therefore, operators rely on batched operations^7^, selecting a threshold for some samples and applying the same threshold to others or optimizing a single threshold simultaneously for multiple samples, resulting in an overall lower F1-score. PhenoBIC still consistently outperforms the sample-level manual gating, demonstrating the benefit of analyzing the information-rich cell biomarker imprint over reducing expression to a single average value. Furthermore, although both annotators exhibit comparable overall performance (Fig. 6A(iv)), the gated outputs vary significantly on a sample-to-sample basis (Fig. 5C, Fig. 6B). As PhenoBIC is an entirely automated, pre-trained deep learning model, it eliminates operator-induced subjectivity.

Additionally, PhenoBIC maintains a minimal difference between precision and recall, reducing the potential bias in subsequent analyses based on cell phenotypic abundance and spatial distribution. The composite precision and recall for the internal and external validation datasets are shown in Fig. 5A(i) and Fig. 6A(i), respectively. Extended Data Figs. 3, 4, and 5 further illustrate the precision and recall balance for the individual cases of the internal and external validation datasets.

We observed that errors by PhenoBIC tended to manifest in specific failure modes that are well-characterized confounders inherent to multiplexed imaging. False positives predominantly occurred in cells exhibiting spatial bleed-through of signals from adjacent cells that were genuinely positive (Extended Data Fig. 6A(iii)), as well as in cells located within regions characterized by diffuse, non-localized, and/or non-specific staining (Extended Data Fig. 6A(iv)). In both cases, the local imaging context was the primary driver of misclassification, and incorporating a wider neighborhood context into the cell-crop representation could further enhance the performance in future iterations. Conversely, false negatives were identified in cells that, although not as brightly stained as others within the region, still demonstrated weak yet specific and localized marker patterns. Next, we sought to quantify the amount of error arising from each of these situations.

We selected five additional held-out patient WSIs and annotated cells specifically in the regions that experience the two types of staining that would potentially lead to false positives: spatial bleed-through (Extended Data Fig. 6A(iii)) and non-specific staining (Extended Data Fig. 6A(iv)). We also annotated typical, non-edge case cells of clear marker-positive (Extended Data Fig. 6A(i)) and marker-negative (Extended Data Fig. 6A(ii)) staining. To isolate the contribution of each failure mode independently, we performed targeted labelling of these four cases in the five WSIs across CD3, CD45RO, and CD8 marker channels (n=9,578 cell labels) to compile the dataset shown in Extended Data Fig. 6E. The PhenoBIC model trained with a binary cross-entropy loss function exhibited poor accuracy in the two false positive failure modes (Extended Data Fig. 6E(i)). Conversely, using a dice loss function instead improved the performance in the false positive failure modes with a minimal increase in false negatives (Extended Data Fig. 6E(ii)). This is mechanistically intuitive, as binary cross-entropy loss optimizes overall accuracy and can under-penalize edge cases, whereas dice loss is robust to this. The prevalence of edge cases within a sample and the accuracy achieved in these instances fundamentally influence the overall performance of the model across the entire sample. For example, varying degrees of non-specific staining can occur due to an array of technical and biological factors. The count of cells exhibiting spatial signal spillover is contingent upon the density of marker-positive cells in proximity to marker-negative cells. Extended Data Fig. 6F illustrates the improvement in performance resulting from the application of a dice loss function to fully annotated samples of the in-house internal validation slides.

### PhenoBIC: Data preprocessing, computational efficiency, and community deployment

Data preprocessing is a crucial part of deep learning inference and can significantly affect model performance, necessitating the selection of a reliable strategy in a data-driven manner. This is conventionally done by percentile-based clipping using the full channel image, compressing the dynamic range of the true positive signal. To address this fundamental limitation, we devised a clipping strategy exclusively utilizing segmented cell regions for optimal contrast enhancement and classification performance (Extended Data Fig. 7, see Methods). We observed stable PhenoBIC performance across our validation datasets using either FOV-level or WSI-level normalization strategies (Extended Data Fig. 7, see Methods) because both were intentionally incorporated in the training data. In contrast, other deep learning algorithms exhibited significantly sensitive performance to normalization at the FOV- versus slide-level. For TMAs, we empirically determined that core-level normalization yields slightly better performance than slide-level normalization (Extended Data Fig. 8, see Methods) To assess the computational efficiency of the PhenoBIC pipeline, we measured the run times for three channels of the in-house internal validation WSIs. PhenoBIC demonstrated ∼3.6x and ∼3.8x speed increases over Nimbus (state-of-the-art model for pre-trained automated cell phenotyping) with CPU and GPU use during deep learning inference, respectively (Fig. 5B). This improvement is attributable to our use of a lightweight convolutional neural network with an EfficientNetV2-S architecture, coupled with the implementation of multi-threading for parallel processing of cells to generate the crops of cell marker imprints prior to deep learning inference. Despite being lightweight, we show that EfficientNetV2-S has adequate model complexity for robust phenotyping because it matched and, in some cases, outperformed an EfficientNetV2-L model (with more trainable parameters) trained with the same dataset and training procedure (Fig. 5A(i) vs 5A(ii) and Fig. 6A(i) vs 6A(ii)). Additionally, we found ∼2.5x average reduction in the total runtime of PhenoBIC when utilizing our GPU-enabled system in comparison to the CPU-only system (Fig. 5B). PhenoBIC can process ∼15,000 cells per minute per channel using CPU only and ∼35,000 cells using our GPU-enabled system. The deep learning inference during runtime exhibits a linear relationship with the number of cells, as illustrated in Extended Data Fig. 2B. The duration of the PhenoBIC inference remains unaffected by the size of the image, relying solely on the number of cells and channels.

We have made the in-house curated datasets for training and testing (https://huggingface.co/datasets/mittal-research-lab/PhenoBIC-Data), the PhenoBIC model (https://huggingface.co/mittal-research-lab/PhenoBIC), and the code to run the pipeline (https://github.com/Shachi-Mittal-Lab/PhenoBIC) open-source for community use. This encompasses approximately 1.4 million curated ground truth cell expression labels alongside the fully trained model weights and the inference pipeline, providing the community with both a deployable tool and a high-quality benchmark resource for future method development. PhenoBIC can be run directly in Python by inputting FOVs of multiplexed data and associated cell segmentation masks or by inputting the multiplex WSI as a single file and associated cell segmentation bounding box coordinates in tabular form.

To enable widespread adoption without direct coding in Python, we have developed an extension (https://github.com/Shachi-Mittal-Lab/PhenoBIC-qupath-extension) that can be deployed in QuPath’s^37^ user interface for immediate generation, visualization, and QC of marker positivity predictions. QuPath is the most widely adopted open-source platform for multiplexed image analysis, positioning PhenoBIC as directly accessible to the large existing QuPath user base, including spatial biology researchers, pathologists, and translational scientists.

## Discussion

In this work, we developed a pre-trained model for automated cell phenotyping of multiplexed data with four key contributions: high classification performance, the largest publicly available high-quality training data spanning millions of annotations, computational efficiency, and community accessibility. Our first contribution is an automated, operator-free cell marker positivity classifier model with robust performance across diverse imaging contexts (tissue types, marker channels, imaging platforms, imaging institutions, cell segmentation algorithms, etc.). We achieved this by combining intentional diversity in our training set with a state-of-the-art convolutional neural network architecture to learn deep representations of cell biomarker imprints. Benchmarking on independent validation datasets shows improvement over manual and other pre-trained approaches in both F1-score and the disparity between precision and recall to avoid systematic downstream bias in cell phenotype frequencies and spatial analyses. We show robust zero-shot performance on imaging platforms and marker channels absent from training. Our second contribution is the largest dataset to-date of paired multiplexed data and manually curated, quality-controlled marker positivity labels. We used these datasets for the training and internal validation datasets in this study and have made them publicly available. These are intended to serve as a resource for the community to develop and evaluate other cell phenotyping models, or for completely different purposes where labeled cell marker positivity is of interest. Our third contribution is PhenoBIC’s computational efficiency, significantly reducing the cell phenotyping runtime within the multiplexed image analysis pipeline. Our fourth contribution is community accessibility via Python code or as an extension utilizing the QuPath interface, enabling use without programming expertise. In particular, PhenoBIC is useful for a priori known cell types, where the biological cell phenotype for specific marker positivity co-expressions are known at the time of study (e.g., cytotoxic T cell = CD3+CD8+ cell).

We show that PhenoBIC outperforms manual gating, which was used as a proxy for methods that use integrated expression as opposed to directly reading image data. In the Nimbus paper^33^, they show that they outperform other computational methods that use integrated expression (e.g., STELLAR and MAPS). Here, we demonstrate that PhenoBIC improved upon the state-of-the-art tool, Nimbus, with respect to F1-score, the gap between precision and recall, and computational efficiency. The public release of the Pan-M dataset^33^ was instrumental in expanding training and validation data diversity in this work. The availability of both the Pan-M dataset and the PhenoBIC dataset will act as powerful complementary tools for the community.

Despite its strong overall performance, PhenoBIC has some limitations that warrant consideration and motivate future development. It underperforms for inherently diffuse markers whose signal interpretation depends on the surrounding tissue architecture (e.g., PD-1 (Fig. 5D(ii)) and CD57 (Fig. 5D(iii)). We also observed that in some large, irregular clusters of under-segmented cells (more than five touching cells erroneously combined as a single cell) in the public datasets, the model had difficulty identifying marker positivity, even if some cells within the cluster showed specific expression. This is because the bounding box of the aggregate cluster is considerably large, leading the model to consider the specific signals as noise within a single cell’s context. Future iterations of PhenoBIC would benefit from incorporating multi-scale or spatially aware input representations that encode the local tissue microenvironment alongside the individual cell imprint. For regions currently experiencing these, we recommend re-training the PhenoBIC model with region-specific examples.

Future work will also focus on expanding training data diversity to include a broader range of diffuse markers, tissue types, and imaging platforms. In line with our goal of promoting community adoption, we will continue developing the pipeline that allows clinicians and researchers to easily integrate PhenoBIC, such as rapid QC through spot-checks and real-time correction in areas where PhenoBIC’s zero-shot performance is sub-optimal. PhenoBIC represents a pathway to generate a spatial cellular landscape of tissue across dozens of markers spanning millions of cells in an objective, reproducible, and high-throughput manner. This can potentially transform multiplexed imaging from a specialized research discovery tool to a scalable technique for both fundamental discovery and clinical translation.

## Methods

### Curating the PhenoBIC Datasets

#### Training & internal validation in-house WSIs

To generate the in-house PhenoBIC training dataset, we acquired and annotated multiplexed immunofluorescence (mIF)-stained images of breast cancer surgically resected formalin-fixed paraffin-embedded (FFPE) sections (n=43 whole slide sections). The tissue sections were stained with the Opal™ 9-color workflow (Quanterix). The biomarker panel and respective fluorophores are as follows: CD206 (Opal 480), CD45RO (Opal 520), CD14 (Opal 540), CD3 (Opal 570), CD163+CD68 cocktail (Opal 620), CD8 (Opal 650), CD66b (Opal 690), and CD4 (Opal 780). Nuclear counterstaining was performed with DAPI. Digital multispectral fields were acquired using the PhenoImager™ HT (Quanterix), previously referred to as Vectra Polaris (Akoya Biosciences), imaging system and stitched to produce contiguous WSIs. The multispectral WSIs were spectrally unmixed using inForm® tissue analysis software (Quanterix).

Out of the 43 mIF-stained breast cancer patient WSIs, three slides were used to annotate CD8 cell labels for PhenoBIC training. Three additional WSIs were used for internal validation, to evaluate PhenoBIC’s cell phenotyping in the same patient cohort, using the same staining and imaging procedures as during training to provide an upper bound for the performance. The six slides were annotated for CD8 cell expression status. First, an expert human annotator selected ∼30-50 cells in each WSI and annotated them as CD8+ or CD8- in QuPath. Then, we trained a rudimentary machine learning model in QuPath with a multilayer perceptron (MLP) neural network using 20 cell intensity data features. These features included the mean, median, standard deviation, minimum, and maximum CD8 signal pixel intensity within the nucleus, cytoplasm, cell membrane, and whole-cell compartment masks. The six trained MLP classifiers were applied to the cells segmented across their respective WSIs for scalable, albeit noisy, annotation across the six slides (Fig. 2A(iv)). Next, rectangular boxes were drawn across the WSIs (Fig. 2A(i)), and a second human annotator manually performed a quality control (QC) review of each rectangular field of view (FOV) and manually overturned CD8 labels where necessary (Fig. 2A(v)). Lastly, a third human annotator performed a second round of QC and noted cells that they thought were still mislabelled and discussed these discrepancies with the second annotator to reach a consensus label for the ambiguous cells (Fig. 2A(vi)). The annotation workflow is shown in Fig. 2A, and the resulting datasets are shown in Fig. 2A(vi).

Targeted labeling was done in five other patient slides from this dataset. Cells were annotated that fell into four categories: CD8+ cells (Extended Data Fig. 6A(i)), “typical” CD8- cells with minimal signal within the cell pixels (Extended Data Fig. 6A(ii)), CD8- cells that experienced signal spillover from neighboring CD8+ cells (Extended Data Fig. 6A(iii)), and CD8- cells in regions of non-specific staining (Extended Data Fig. 6A(iv)). Similarly, cells were also annotated for CD45RO and CD3 expression status to evaluate phenotyping performance across the four cell stain categories to yield the dataset shown in Extended Data Fig. 6E. The objective of this targeted label dataset was to assess the performance of PhenoBIC in two typical cases and two edge cases, which generally represented the main failure modes of the model.

#### Training & internal validation Pan-M FOVs

To supplement our training and internal validation datasets, we utilized Pan-M^33^, a public dataset of multiplex images with paired cell expression labels across various marker channels. Specifically, we used the part of the dataset with paired “gold-standard” annotations, which refer to cell expression labels that were manually proofread, to minimize noise in the ground truth labels. We used three subsets of the dataset (with three labelled FOVs in each): colon tissue imaged using Vectra Polaris, pancreas tissue imaged using Vectra Polaris, and decidua tissue imaged using Multiplexed Ion Beam Imaging (MIBI). One labelled FOV from each dataset was added to the training data pool, while the remaining two were used for internal validation. From the labelled datasets, marker channels with no positively labeled cells in an FOV were excluded from the training and validation processes.

#### External validation TMAs

We utilized three TMAs annotated with cell expression labels for various marker channels to evaluate the generalizability of PhenoBIC. Since the patient cohorts, marker panels, staining, and imaging runs differ from the training dataset, these TMAs serve as external validation datasets. The number of cores, marker channels, and cells annotated is shown in Extended Data Fig. 1B. The annotation workflow is shown in Fig. 2B. Unlike the training and internal validation WSIs, we found that a single MLP classifier was not effective for scalable cell marker positivity annotation across different cores within a TMA. Since the cores originated from different patient samples and were processed separately, there was a systematic variation between cores. Consequently, we adopted a more direct annotation method, demonstrated using a representative TMA core’s FOV for the CD45RO marker channel shown in Fig. 2B(ii)-(v), where the entire core was divided into rows and columns, and individual tiles were annotated for marker-positive cells. Each tile was annotated during the same session, and well-defined resting time was incorporated between sessions to prevent mislabeling due to annotator exhaustion. Once all the positive cells were marked, marker-negative cells were computationally identified as unlabeled cells. A second human annotator conducted a quality control (QC) review of each TMA core FOV and discussed any discrepancies with the first annotator to reach consensus on cell labels. In this way, we annotated cores sampled from three TMAs for cell expression labels of various markers to construct the external validation dataset for PhenoBIC evaluation.

The breast cancer tissue sections in TMA #1 were stained using the Opal™ 7-color workflow (Quanterix). The biomarker panel and corresponding fluorophores are as follows: CD3 (Opal 480), FoxP3 (Opal 520), CD45RO (Opal 570), CK (Opal 620), CD4 (Opal 690), and CD8 (Opal 780). Whole slide scans were captured using the PhenoImager™ HT (Quanterix) imaging system. The multispectral WSIs were spectrally unmixed with inForm® tissue analysis software (Quanterix). TMA #2 contains cores from the same samples as TMA #1 and was stained and imaged with the same protocol. However, the mIF panel used was different: CD8 (Opal 480), PD-L1 (Opal 520), PD- 1 (Opal 570), CD163+CD68 cocktail (Opal 620), CD20 (Opal 690), and CD4 (Opal 780). TMA #3 contains breast cancer tissue cores with different samples from TMAs #1 and #2. The following PhenoCycler® (Quanterix), formerly known as CODEX (Akoya Biosciences), panel was run: HLA-DR, Ki67, CD3e, CD8, CD14, IDO-1, CD4, CXCR2, FoxP3, CD141, CD20, LEF-1, CD138, CD15, E-cadherin, PD1, MHC I, PDL1, CD11b, CD44, CD34, IFN***γ***, Vimentin, CD68, LAG-3, CK, CD11c, CD163, CD79a, CD66b, VISTA, Granzyme B, CD107a, CD38, CXCR3, CD27, CD56, TOX, EpCAM, Podoplanin, CD45, CD45RO, Collagen IV, PCNA, Galectin-3, MPO, aSMA, and TCF7. Nuclear counterstaining for all three TMAs was performed with DAPI. Cores with no or minimal artifacts, such as tissue tearing, tissue folding, and out-of-focus, were considered for sampling to be annotated as ground truth. Cores were chosen to maximize the geographic distribution of the sampling across the TMAs, where the sampling was blinded to PhenoBIC outputs for unbiased evaluation of the performance.

### PhenoBIC pipeline

The input to the classifier is a crop of the buffered bounding box of a cell’s mask of the biomarker channel of interest (Fig. 1E(i)-(iv)). The bounding box of a cell is buffered by 10% in all directions (i.e., 20% longer and wider than the original bounding box) to include the immediate pixels neighboring the cell as input to the classifier. This provides some context to the classifier about the signal from regions adjacent to the cell. Our developed pipeline includes functionality that allows users to set any amount of buffering they prefer. The buffered bounding box is cropped, pre-processed, and resized to 48×48 pixels using nearest-neighbor interpolation.

For efficiency, we utilize parallel computing threads for the generation of cell crop batches, subsequently performing PhenoBIC inference on the preprocessed cell crops, which leverages GPU computation if available. PhenoBIC makes independent and sequential inferences across the multiplex channels to predict the cell expression (+ vs -) for each marker. Cell phenotypes can then be assigned based on a priori known combinations of marker expression. Our digital pipeline can be run on a batch of multiplex images so that nuclear segmentation and cell phenotyping happen sequentially for each sample, enabling automated and operator-independent single-cell spatial profiling.

### Cell segmentation

The PhenoBIC pipeline accepts two inputs: multiplexed images and cell segmentations. These cell segmentations may be supplied to our pipeline either as mask files or as cell bounding box coordinates in a tabular format. To enhance the heterogeneity of our training and validation datasets, we employed cell segmentations derived from two distinct approaches. Nuclear segmentations for our in-house datasets were produced utilizing an in-house fine-tuned Cellpose model using the previously described approaches^41^. Subsequently, whole-cell boundaries were approximated by buffering the convex hull of the nucleus prediction masks by 5 μm, with a maximum expansion distance limited to 1.5 times the nucleus region about its centroid, and overlaps between cells were resolved through Voronoi tessellations^42^. This whole-cell approximation process from nuclear predictions is a common approach used in previous studies^43–47^. For the Pan-M public datasets, whole-cell segmentation was carried out using the Mesmer^11^ algorithm^33^.

### Preprocessing

For each marker channel, we clip the signal using upper and lower normalization constants to handle outlier pixels, then normalize the image by linearly scaling between 0 and 1. Following clipping and normalization, the normalized pixel intensities within the cell bounding box are multiplied by 255 and converted to integer values. The design of this clipping strategy is non-trivial, as marker-positive cells are typically a sparse minority of the total cell population, where the conventional percentile-based clipping applied to the full channel image can compress the dynamic range of the true positive signal. Initially, we tested preprocessing without any clipping (Extended Data Fig. 7A), and then with clipping at 0.01-99.99 percentiles of each channel (Extended Data Fig. 7B). This clipping approach led to a notable decrease in precision as it would overamplify background signal, making the normalized signal too bright for true negative cells. Therefore, clipping based on a pre-defined percentile of the entire channel image is not a scalable method for preprocessing data for this task.

Our clipping method calculates the 90*th* and 10*th* percentiles for each channel within each cell segmentation region. It then selects the minimum of the lower clip and the maximum of the upper clip across all cells during preprocessing as the normalization points for the image channel intensities. We observe that this method does not result in a dramatic decrease in precision; rather, it improves the F1-score and significantly reduces the difference between precision and recall (Extended Data Fig. 7D). We also experimented with less clipping (0*th*-100*th* percentile) and more clipping (20*th*-80*th* percentile) but found these to be less optimal (Extended Data Figs. 7C and 7E).

In addition to the clipping strategy, the size of the preprocessing field is also a significant factor contributing to the model inference. For example, users may acquire images in a targeted manner only for FOVs of interest and input these into the model, or alternatively, they could provide an entire multiplex WSI to the model. Each approach results in different clipping outcomes and variably normalized inputs to the model. To account for this heterogeneity during model training (Fig. 1G), our in-house annotated FOVs were preprocessed using clipping parameters derived from the entire slide (slide-level preprocessing), whereas the public Pan-M FOVs were preprocessed using clipping parameters obtained for each FOV individually (FOV-level preprocessing). To assess the generalizability of this approach for future users, the in-house internal validation FOVs (Fig. 5C, Extended Data Fig. 7(i)) were preprocessed using a slide-level approach, while the Pan-M internal validation FOVs (Fig. 5D, Extended Data Fig. 7(iii)) were preprocessed at the FOV level. Similarly, the external validation FOVs (Fig. 6, Extended Data Fig. 7(ii)) were preprocessed at the FOV level, so that each core is normalized independently. With both approaches, FOV and WSI-level preprocessing, we show robust performance by PhenoBIC. At the TMA level, we further demonstrate that even though there is wide variation in the dynamic range across cores (Extended Data Fig. 8A), it does not significantly impact the F1-score using either the core level or slide level normalization (Extended Data Fig. 8B). This provides evidence that PhenoBIC’s classification is primarily driven by the spatial morphology of the staining pattern within a cell crop rather than just the absolute intensity values, representing a true biomarker imprint. Overall, we recommend that future users employ core-level preprocessing as it is accompanied by a reduced disparity between precision and recall (Extended Data Fig. 8B).

### PhenoBIC model architecture

We utilize a convolutional neural network (CNN) with an EfficientNetV2-S^48^ architecture as PhenoBIC’s feature extraction backbone. The input image (48×48×3 pixels) represents the marker channel crop of a cell with the biomarker imprint pattern. A global average pooling layer is applied to the output feature map, collapsing the spatial dimensions and generating a 1,280-dimensional feature vector for each input image. This pooled feature vector is then passed through a shallow fully-connected classification head, which comprises an intermediate dense layer with 512 neurons and a final sigmoid-activated neuron for binary output prediction (marker-positive/marker-negative). The complete model comprises ∼21 million parameters. We also trained an experimental model with a much larger feature extraction backbone (EfficientNetV2-L^48^) and the same classification head design (resulting in ∼118 million parameters), but it didn’t show any improvements over EfficientNetV2-S (Fig. 5A(ii), Fig. 6A(ii)).

### Training details

#### Dice loss models

We employed a dice loss function, which is mathematically associated with the F1-score, for training both the experimental EfficientNetV2-L and the final EfficientNetV2-S PhenoBIC model, in order to align the training objective with the optimal evaluation criteria of minimizing both false positive and false negative predictions. We implemented a four-step training approach. Initially, the feature extraction backbone with pre-trained weights from ImageNet was loaded. Subsequently, the classification head, comprising an intermediate dense layer and an output dense layer with randomized weights, was added atop the feature extraction block. During the first training round, only the weights of the classification head were trained using a binary cross-entropy loss function with a high learning rate of 10^-4^. In the second training round, all model weights, including the feature extraction block, were trained with binary cross-entropy loss and a reduced learning rate of 10^-6^. The third round involved training only the classification head weights with a dice loss function with a higher learning rate of 10^-4^. In the final phase, all model weights, inclusive of the feature extraction block, were trained with dice loss and a lower learning rate of 10^-6^. This tiered training strategy of initially optimizing only the classification head, followed by fine-tuning of all network weights, and transitioning from binary cross-entropy to dice loss was employed to promote stable training and improved convergence.

During the training steps, an Adam optimizer was employed with a batch size of 32. A dropout rate of 0.2 was implemented on the intermediate dense layer subsequent to feature extraction, introducing random zeroing of 20% of the activations during training to reduce overfitting prior to the final classification layer. To enhance generalization and stability during training, data augmentation techniques were applied to the training set, including random rotations up to 40 degrees, as well as random horizontal and vertical flips of the cell crops containing the biomarker imprints. The training was monitored using a validation subset (30% data split) of the training dataset. Early stopping was utilized based on performance metrics on the validation subset to prevent overfitting, complemented by a learning rate reduction strategy triggered by stagnation in performance. During the initial two rounds, which employed binary cross-entropy loss, validation accuracy was monitored; in subsequent rounds utilizing dice loss, validation F1-score was monitored at the end of each epoch.

#### Binary cross-entropy loss model

To highlight the impact of employing a dice loss function, we trained an EfficientNetV2-S model utilizing a binary cross-entropy loss function for comparative purposes. A two-phase training approach was adopted, wherein the initial phase involved training solely the classification head weights with a higher learning rate, followed by a second phase that entailed training all model weights with a lower learning rate. A comparable strategy was applied regarding the optimizer, batch size, data augmentation, and validation subset accuracy monitoring as was employed with the dice loss models. Given that binary cross-entropy loss is vulnerable to class imbalance, we undersampled the cases labeled as marker-negative and oversampled those labeled as marker-positive (employing data augmentation) to create a class-balanced training dataset. Consequently, we demonstrated that the dice loss function enhanced performance in failure modes relative to binary cross-entropy loss (Extended Data Fig. 6E), even after addressing the class imbalance that binary cross-entropy loss is known to struggle with.

### Other phenotyping tools for comparison

#### Manual gating

Two trained users, “Annotator #1” and “Annotator #2,” performed cell expression classification through manual gating. They manually selected thresholds based on the mean measurements of the marker channels (average pixel intensity of biomarker signal within the cell region) by visualizing in the QuPath interface. Cells with measurements above the threshold were classified as “positive,” while those below were “negative” for the biomarker. For the internal validation WSIs, one gate was selected for each patient WSI (WSI-level gating). For the external validation TMAs, one gate was selected for each core (core-level gating) to account for differences in tissue sample preparation, staining, and biological nuances inherent to different patient samples. The performance of automated and operator-independent PhenoBIC cell phenotyping was compared to the sample-level manual gating as a reference.

#### Nimbus

Nimbus is a recently developed pre-trained deep learning model for cell marker expression classification^33^. We implemented Nimbus by installing the Python package “nimbus-inference” v0.0.5. We used the default parameter inputs in the inference function and a “normalization quantile” of 0.9999, as in their publication^33^. The performance and computational efficiency of PhenoBIC were compared with Nimbus as a benchmark using the validation datasets.

### Evaluation metrics

The performance of PhenoBIC, Nimbus, and manual gating was quantitatively evaluated by comparing them with the ground truth annotated cell marker expression labels across the validation datasets. The number of true positive (TP), false positive (FP), true negative (TN), and false negative (FN) cell predictions was computed. From here, the precision, recall, F1-score, and accuracy metrics were computed as evaluation metrics for cell phenotype classification. Precision measures the proportion of positive predictions that are true positives, while recall assesses the ratio of actual positive cells correctly identified. The F1-score is the harmonic mean of precision and recall, representing the model’s ability to balance false positives and false negatives. Accuracy indicates the total number of cell predictions that were correctly classified. The evaluation metrics can be calculated as follows:

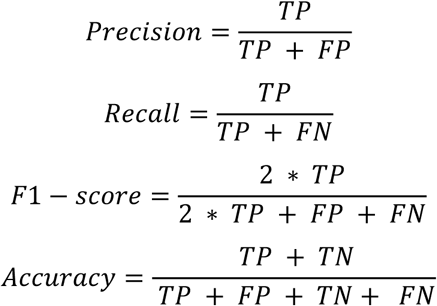

We included the accuracy metric in the composite evaluations (Fig. 5A and Fig. 6A) to highlight its shortcomings as an evaluation metric for this task and emphasize the importance of using precision, recall, and F1-score instead. There is a significant class imbalance, where marker-negative cells are nearly always far more prevalent than their marker-positive counterparts. Therefore, the accuracy metric would not represent the true performance of the model. This discrepancy arises because the majority of cells are “true negatives,” and the specificity of the model (which tends to be very high) ends up predominantly influencing the accuracy measure. In this context, a model that classifies every cell as marker-negative would achieve high accuracy while being completely uninformative for phenotyping and downstream analysis.

Therefore, we rely on Precision, Recall, and F1-score as the main metrics for evaluating the model performance as opposed to accuracy or specificity. We also regard the difference between precision and recall for the validation cases as a crucial metric. A smaller difference indicates a superior capacity to balance false positives and false negatives, thereby preventing bias in subsequent single-cell analyses following cell phenotyping

### Runtime analysis

The total runtimes, including pre-processing, deep learning model inference, and other necessary operations of PhenoBIC and Nimbus, were recorded and compared across three marker channels of the three internal validation WSIs (Fig. 5B). For PhenoBIC, the multiplexed data was supplied as a single OME-TIFF file for each WSI, and the cell segmentation data was provided as a table of cell bounding box coordinates per WSI. For Nimbus, the multiplexed data was supplied as single TIFF files for each channel, and the cell segmentation data was provided as a single mask per WSI. We deliberately refrained from pre-tiling the images to permit the algorithms to execute their own tiling processes on the backend, thereby avoiding confounding read/write speeds of the provided tiles and facilitating a more standardized comparison. Both algorithms require the configuration of a back-end tile size for operation. The default setting for PhenoBIC is 10,000 pixels, whereas for Nimbus, it is 1,024 pixels. Regarding PhenoBIC, it was observed that increasing tile sizes can reduce the overhead associated with image reading and thereby enhance efficiency. Although larger tile sizes than 10,000 pixels are feasible, experimentation was limited to this size due to the potential memory constraints faced by computers with limited RAM (e.g., <32 GB). We recommend users to lower the tile size setting if they run into memory issues. Attempts to operate Nimbus with tile sizes greatly exceeding 1,024 pixels resulted in memory issues; consequently, the default size was maintained. The hardware environment we ran both pipelines in is as follows:

- System: Windows 11 Enterprise
- CPU: Intel(R) Core(TM) Ultra 9 285K (3.70 GHz)
- RAM: 128 GB
- GPU: NVIDIA GeForce RTX 5070, 12 GB
- CUDA: version 13.0

## Data and code availability

PhenoBIC can be run via Python code (https://github.com/Shachi-Mittal-Lab/PhenoBIC) or as an extension in the QuPath interface (https://github.com/Shachi-Mittal-Lab/PhenoBIC-qupath-extension). The publicly released inhouse labelled datasets can be found at https://huggingface.co/datasets/mittal-research-lab/PhenoBIC-Data and the PhenoBIC model can found at https://huggingface.co/mittal-research-lab/PhenoBIC.

## Acknowledgements

We acknowledge funding from the Breast Cancer Research Foundation (S.V), University of Washington Institute of Medical Data Science (A.S.K, S.M, S.C.P), NIH/NCI K99CA293004, and the University of Washington Departments of Chemical Engineering and Lab Medicine & Pathology Startup Grants (S.M). This research was supported by the Experimental Histopathology (RRID:SCR_022612) Shared Resource of the Fred Hutch/University of Washington Cancer Consortium (P30 CA015704) and the Translational Pathology (RRID:SCR_028306) core at Fred Hutch.

## Ethics declarations

The authors declare no competing interests.

## Author contributions

A.S and S.M conceptualized and designed the research. K.S.S, A.S.K, C.I.L, S.C.P, and S.V acquired and curated the in-house imaging data. A.S, C.Z, M.G.H, E.A.C, and L.L.C annotated the datasets for cell marker expression. A.S. developed the PhenoBIC pipeline and trained the PhenoBIC model. A.S and K.S.S performed independent manual gating as a comparison to PhenoBIC. A.S and S.M wrote the manuscript.

