## Extended data for "PhenoBIC: operator-free single-cell spatial phenotyping in multiplex imaging data using deep learning of cell staining patterns"

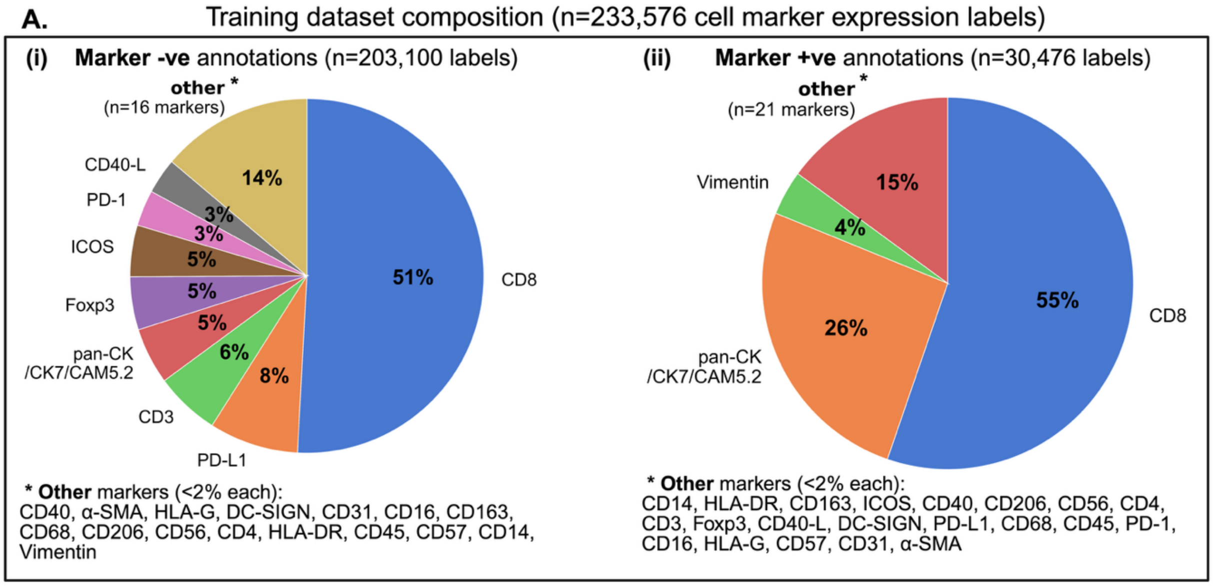


**B.** External validation dataset composition (n=747,551 cell marker expression labels)

**(i) TMA #1**

| Cell marker | Cores annotated | +ve labels | -ve labels |
|---|---|---|---|
| CD8 | 12 | 7,276 | 138,425 |
| CD45RO | 12 | 19,710 | 125,991 |
| CD3 | 2 | 1,171 | 23,293 |
| Foxp3 | 2 | 301 | 24,163 |

**(ii) TMA #2**

| Cell marker | Cores annotated | +ve labels | -ve labels |
|---|---|---|---|
| CD8 | 12 | 6,867 | 151,512 |
| CD20 | 2 | 388 | 23,486 |
| CD163 +CD68 | 2 | 1,373 | 22,501 |
| PD-1 | 2 | 342 | 23,532 |

**(iii) TMA #3**

| Cell marker | Cores annotated | +ve labels | -ve labels |
|---|---|---|---|
| CD8 | 18 | 8,188 | 125,212 |
| PD-L1 | 2 | 51 | 13,783 |
| Pan-CK | 2 | 5,402 | 9,591 |
| Ki67 | 2 | 530 | 14,463 |

- Breast tissue TMAs imaged
- Vectra (TMA #1 & TMA #2) & CODEX (TMA #3) imaging technologies
- Biomarkers tested: **T & B lymphocyte** (CD3, CD8, CD20, CD45RO, Foxp3), **macrophage** (CD163+CD68), **tumor/epithelial** (pan-CK), **proliferation** (Ki67), **immune checkpoint** (PD-1, PD-L1)

**<u>Extended Data Fig.1: Extended breakdown of dataset biomarker composition.</u> A.** Biomarker composition of the training dataset for cell expression **(i)** marker-positive and **(ii)** marker-negative labels. **B.** Biomarker composition and dataset sizes for external validation TMAs **(i)** #1, **(ii)** #2, and **(iii)** #3.

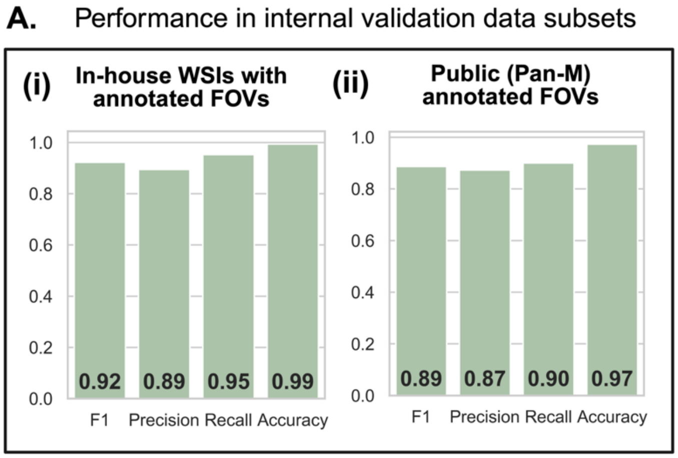


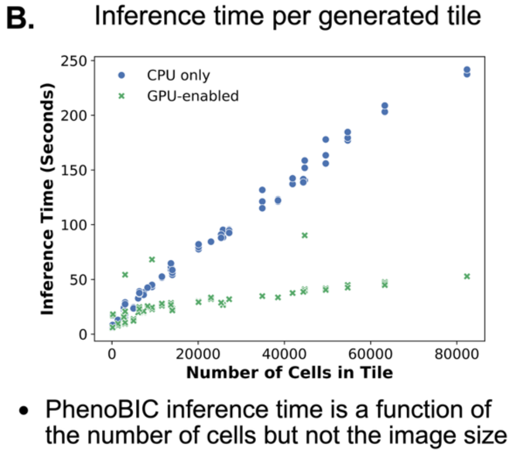

**Extended Data Fig.2: Classifier performance and inference time on internal validation data subsets. A.** Evaluation of PhenoBIC on two subsets of the internal validation dataset: **(i)** In-house multiplex WSIs with FOVs exhaustively annotated for CD8 cell expression (CD8+/CD8-) and **(ii)** a publicly available annotated dataset. **B.** The inference time for each tile generated during the PhenoBIC pipeline as a function of the number of cells. Each data point represents the time taken to run PhenoBIC inference for one tile for one marker channel. PhenoBIC was run on the three in-house internal validation WSIs for three marker channels to generate these time measurements.

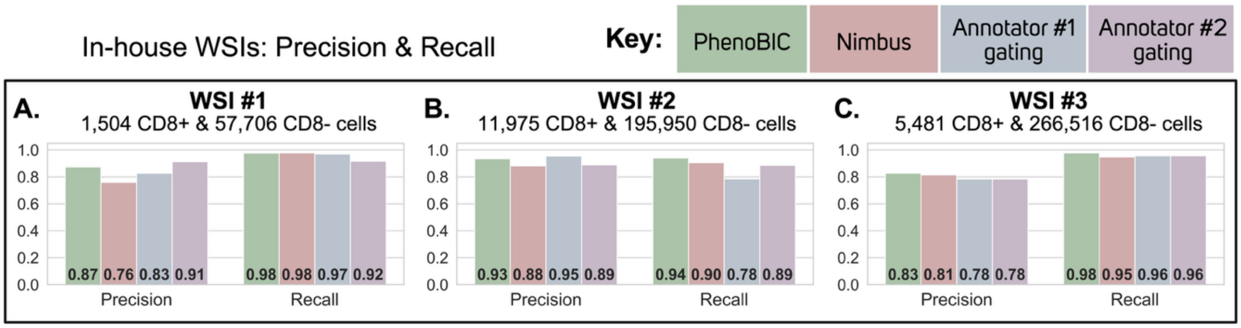


**Extended Data Fig.3: Quantitative evaluation using in-house WSIs of internal validation dataset.** CD8 cell expression classification performance using precision and recall of PhenoBIC outputs, Nimbus outputs, and manual gating by two expert human annotators for the internal validation WSIs **(A)** #1, **(B)** #2, and **(C)** #3.

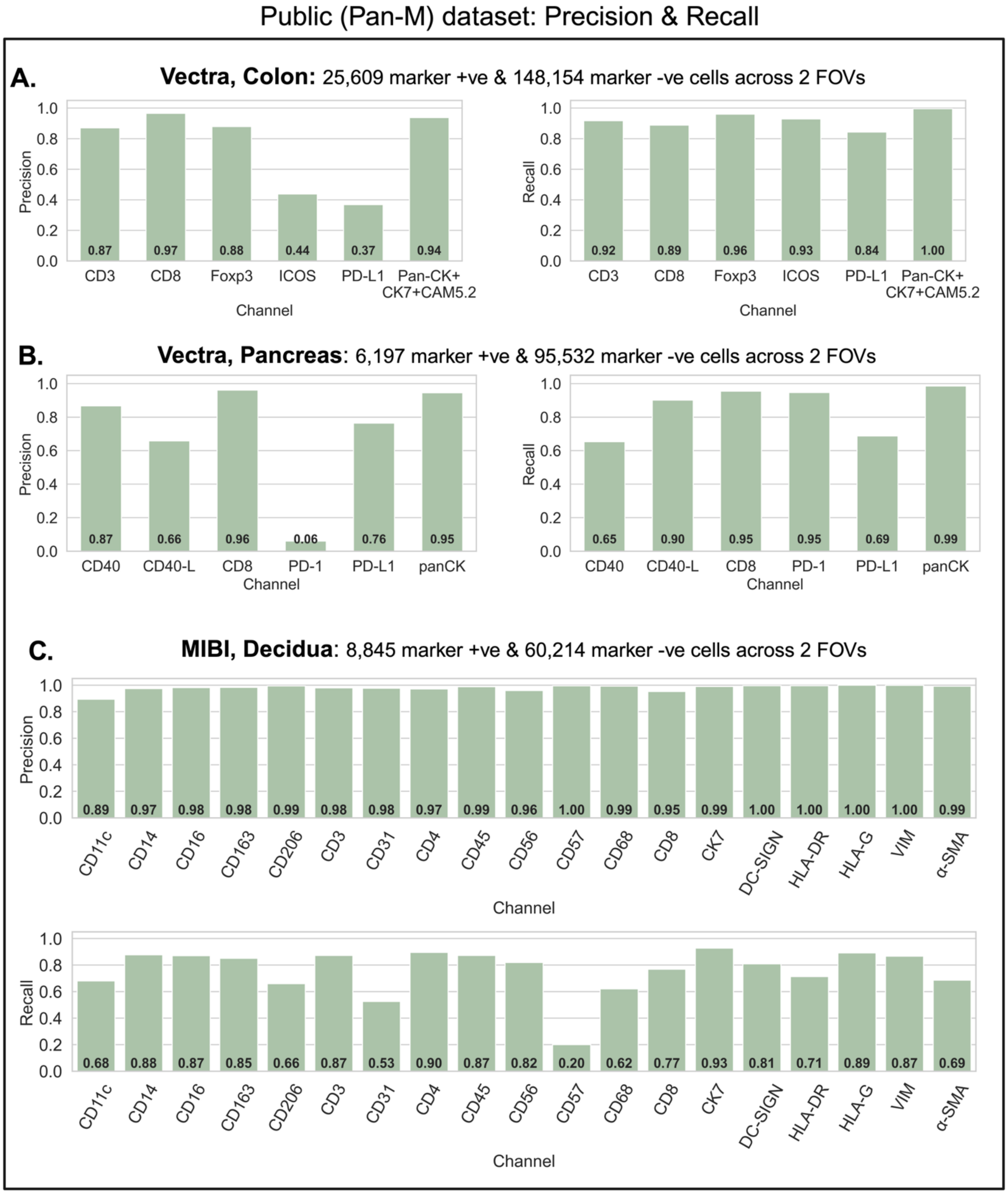


**Extended Data Fig.4: Quantitative evaluation using public (Pan-M) subset of internal validation dataset.** Cell expression classification performance using precision and recall of PhenoBIC for various biomarkers for three subsets of the data with two annotated FOVs in each: **(A)** colon tissue imaged with Vectra, **(B)** pancreas tissue imaged with Vectra, and **(C)** decidua tissue imaged with MIBI.

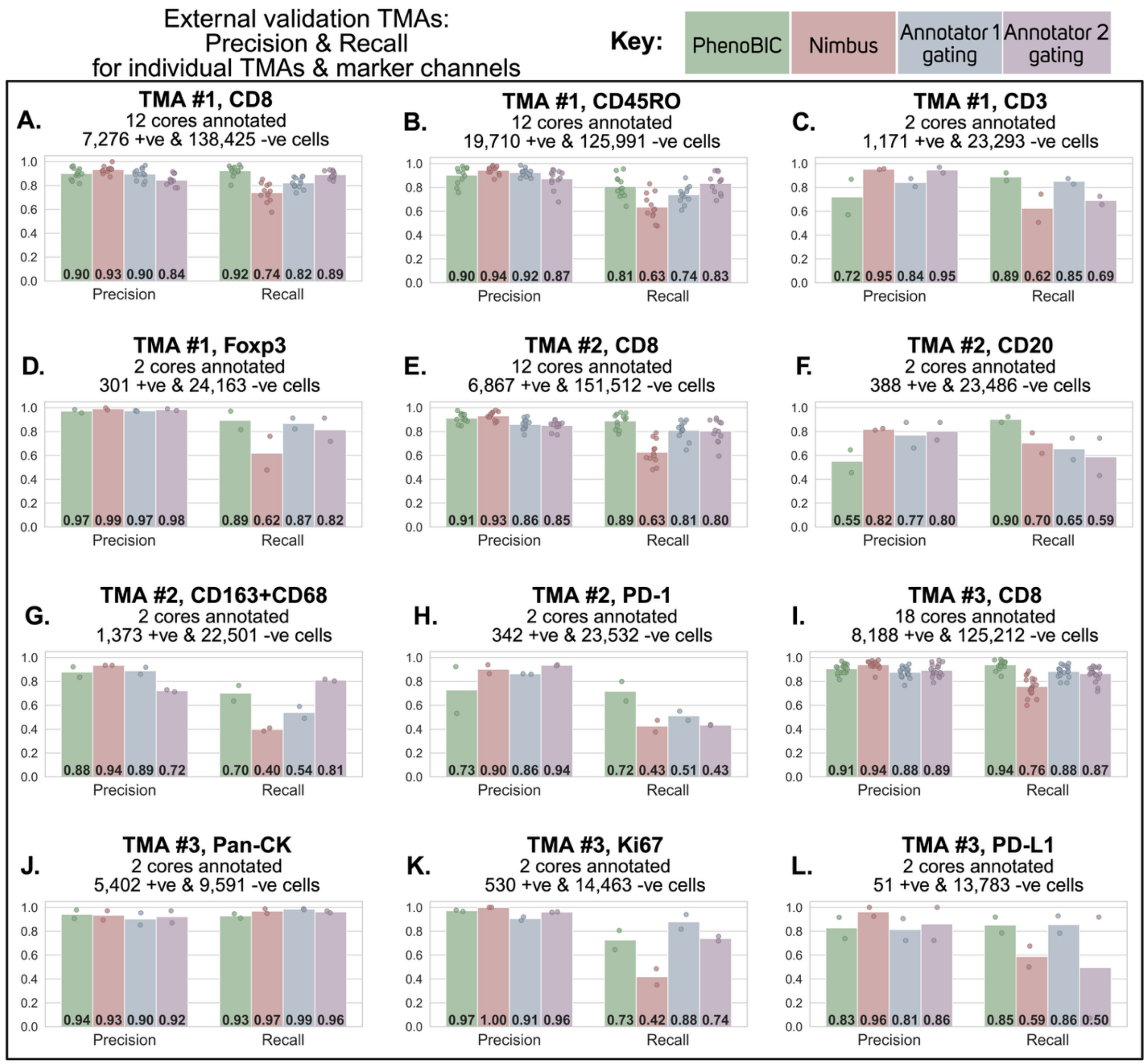


**<u>Extended Data Fig.5: Quantitative evaluation using external validation TMAs.</u>** Cell expression classification performance using precision and recall of PhenoBIC, Nimbus, and manual gating by two expert human annotators for annotated biomarker channels. Results are shown for annotated cores in TMA #1 for channels: **(A)** CD8, **(B)** CD45RO, **(C)** CD3, and **(D)** Foxp3, in TMA #2 for channels: **(E)** CD8, **(F)** CD20, **(G)** CD163+CD68 cocktail, and **(H)** PD-1, and in TMA #3 for channels: **(I)** CD8, **(J)** Pan-CK, **(K)** Ki67, and **(L)** PD-L1. Data points show the precision and recall for individual annotated TMA cores, with the average shown by the bars.

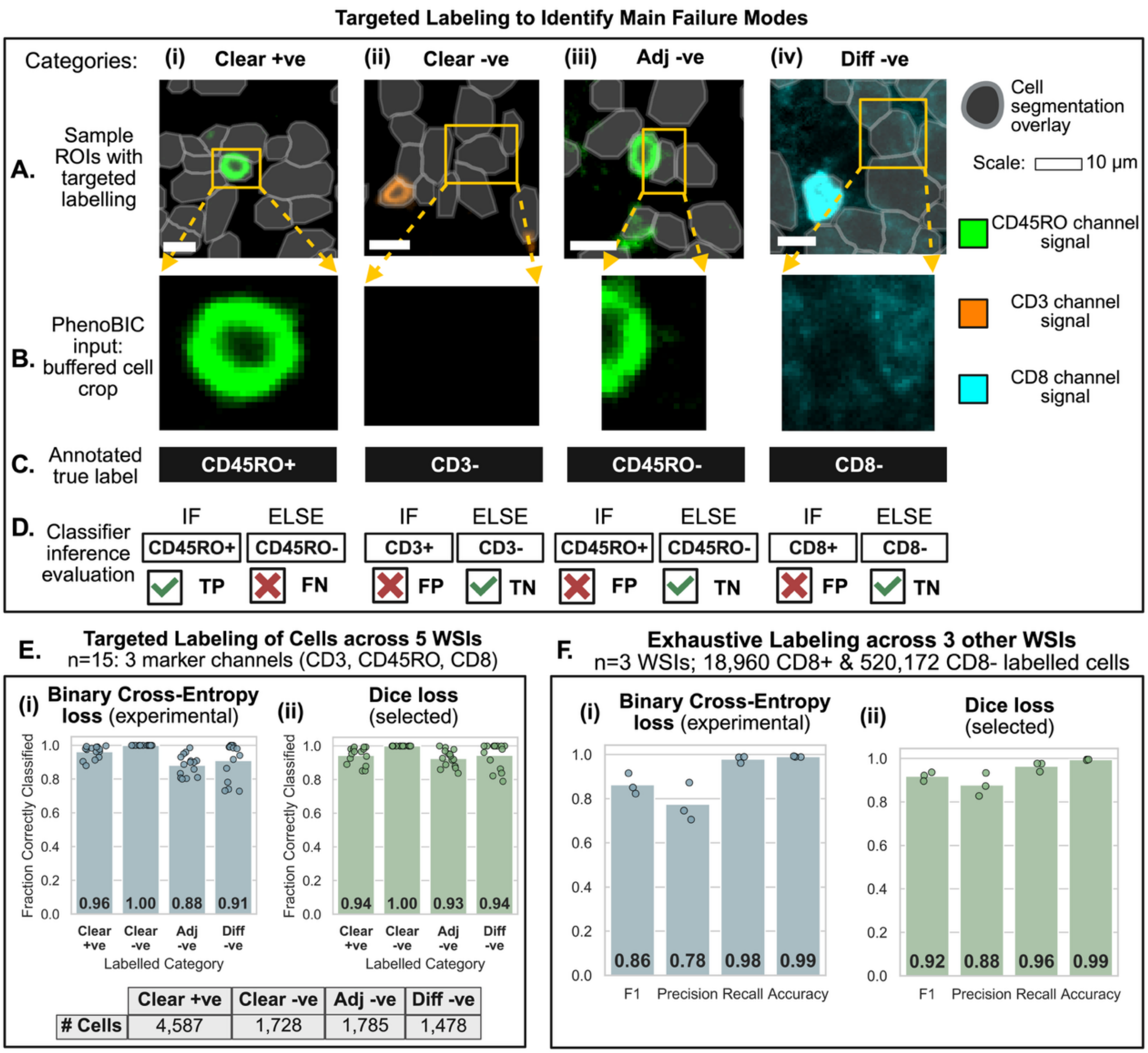


**Extended Data Fig.6: Failure modes.** Cells were annotated in specific regions to interrogate the performance of PhenoBIC in these regions. Five WSIs from the in-house dataset, which were not used for model training, were chosen for this analysis. **A.** Sample regions of interest (ROIs) showing the four categories of labelled examples: **(i)** Clear +ve: cells with typical true-positive staining showing marker expression, **(ii)** Clear -ve: cells with no marker signal showing absence of marker expression, **(iii)** Adj -ve: first edge case where cells that are not expressing the marker have spatial bleed-through of marker signal from an adjacent cell which is marker-positive, **(iv)** Diff -ve: second edge case where cells that should be marker-negative have diffuse, non-localized signals in their region. **B.** Buffered cell crops of the cells annotated across the four label categories. This is the input to the PhenoBIC model. **C.** The annotated label of each cell category. Clear +ve cells were assigned marker "positive" labels, while Clear -ve, Adj -ve, and Diff -ve cells were assigned marker "negative" labels. **D.** PhenoBIC inference classes for the four cell categories can be either True Positive (TP), False Positive (FP), True Negative (TN), or False Negative (FN). The counts of each in a region determine the F1-score, precision, and recall metrics. **E.** The fraction of cells correctly classified across the four cell categories when using a PhenoBIC model trained using **(i)** binary cross-entropy loss (experimental) and **(ii)** dice loss function (selected). **F.** F1-score, precision, recall, and accuracy of PhenoBIC trained using **(i)** binary cross-entropy loss and **(ii)** dice loss, for the internal validation dataset (n=3 WSIs) with exhaustive (i.e., not targeted cell categories) annotations (all cells within chosen FOVs labelled).

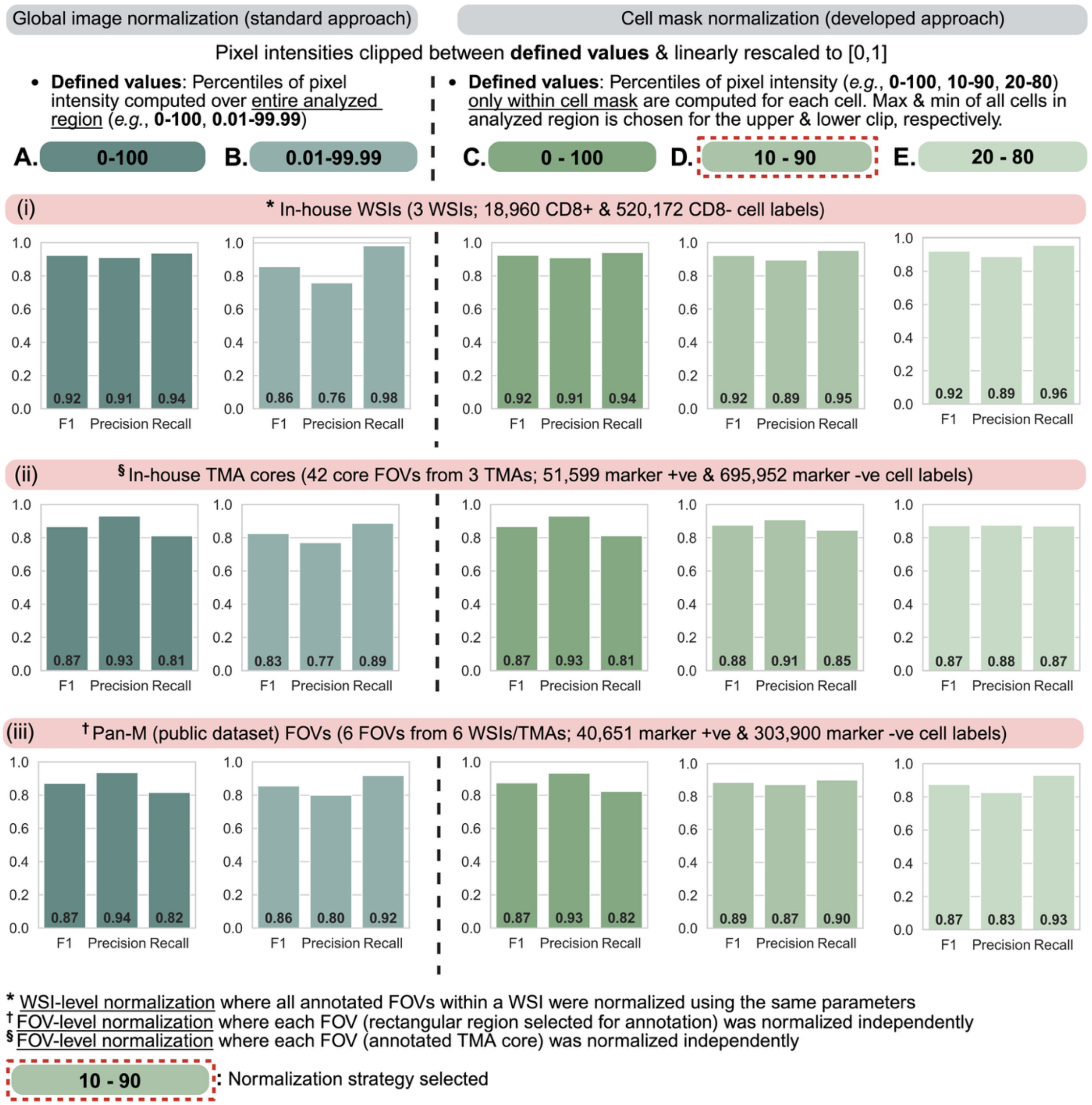


**Extended Data Fig.7: Data preprocessing before PhenoBIC inference.** Two different preprocessing approaches were tried out. **A-B.** Global image normalization is a standard approach in biological image processing, for which two sets of clipping percentiles were investigated. **C-E** Cell mask normalization is the preprocessing approach developed in this work to overcome shortcomings of the global approach, and three sets of clipping percentiles were investigated. 10$^{th}$-90$^{th}$ percentile clipping with the cell mask normalization approach (**D**) was selected as the best strategy. **(i)** FOVs from the internal validation in-house WSIs were normalized at the WSI level, while **(ii)** TMA core FOVs from the external validation TMAs and **(iii)** internal validation FOVs from the Pan-M public dataset were normalized at the FOV level to test both preprocessing field effects.

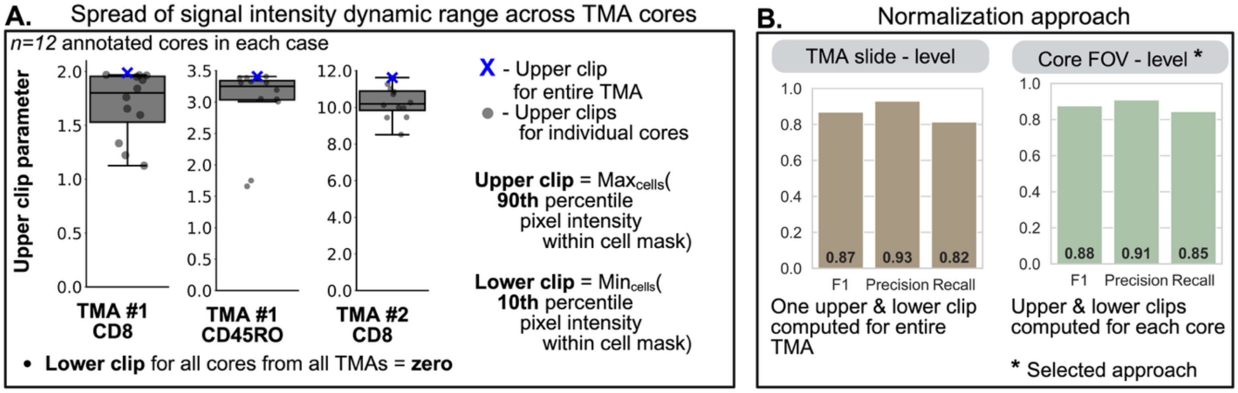


**<u>Extended Data Fig.8: Effect of preprocessing field size for TMAs.</u>** **A.** The spread of the signal intensity dynamic range across cores within a tissue microarray (TMA) slide, as measured between the upper and lower normalization clipping limits (using the developed cell mask normalization approach). Results are shown for the CD8 channels in external validation TMAs #1 and #2 and for the CD45RO channel in TMA #1. The box plots show the upper clips for each core field of view (FOV) as the preprocessing field, while the lower clips were always equal to zero. The upper clip for the entire TMA slide, as the preprocessing field, is marked with a blue cross. **B.** Both preprocessing fields, TMA slide-level and core FOV-level, were investigated prior to inference with PhenoBIC. The core FOV-level showed a marginal performance gain and was selected as the best preprocessing field strategy.
